# G-quadruplex folding uncouples cis-activation from collateral trans-cleavage by Cas12a

**DOI:** 10.64898/2026.07.23.740342

**Authors:** Irina V. Safenkova, Maria V. Kamionskaya, Alexey V. Samokhvalov, Anatoly V. Zherdev, Boris B. Dzantiev

## Abstract

The CRISPR/Cas12a system is widely used in nucleic-acid diagnostics because Cas12a couples guide-directed recognition of a *cis*-target with indiscriminate collateral *trans*-cleavage of ssDNA. Controlling these activities through DNA structure can tune diagnostic signals. G-quadruplexes (G4s) are strong structural regulators, yet their ability to modulate Cas12a activation and collateral cleavage remains undefined. Here we focused on the G4 scaffold (TGGG)n and tested it with LbCas12a and AsCas12a by real-time cleavage assays, circular dichroism, FRET and denaturing PAGE. In the *cis* position, compact (TGGG)n G4s activated Cas12a; for the most stable (TGGG)_5_, k_cat_/K_M_ was 5.9×10^4^ and 8.9×10^4^ M^−1^ s^−1^ for LbCas12a and AsCas12a, respectively. K^+^ reduced *cis*-activation rates by approximately 10-fold, depending on scaffold and temperature. FRET and denaturing PAGE showed that productive *cis*-recognition involves G4 unfolding followed by target-strand cleavage. In the *trans* position, compact G4s fully resisted collateral reporter cleavage. Core disruption and G-rich non-G4 controls restored reporter cleavage, showing that resistance depends on G4 architecture rather than guanine content. These data define a compact G4 scaffold that remains functionally *trans*-resistant while retaining guide-dependent *cis*-target competence. Thus, compact G4 folding provides a programmable structural mechanism for separating Cas12a activation from reporter cleavage, opening a route to signal-gated diagnostic designs.

**Graphical abstract:** 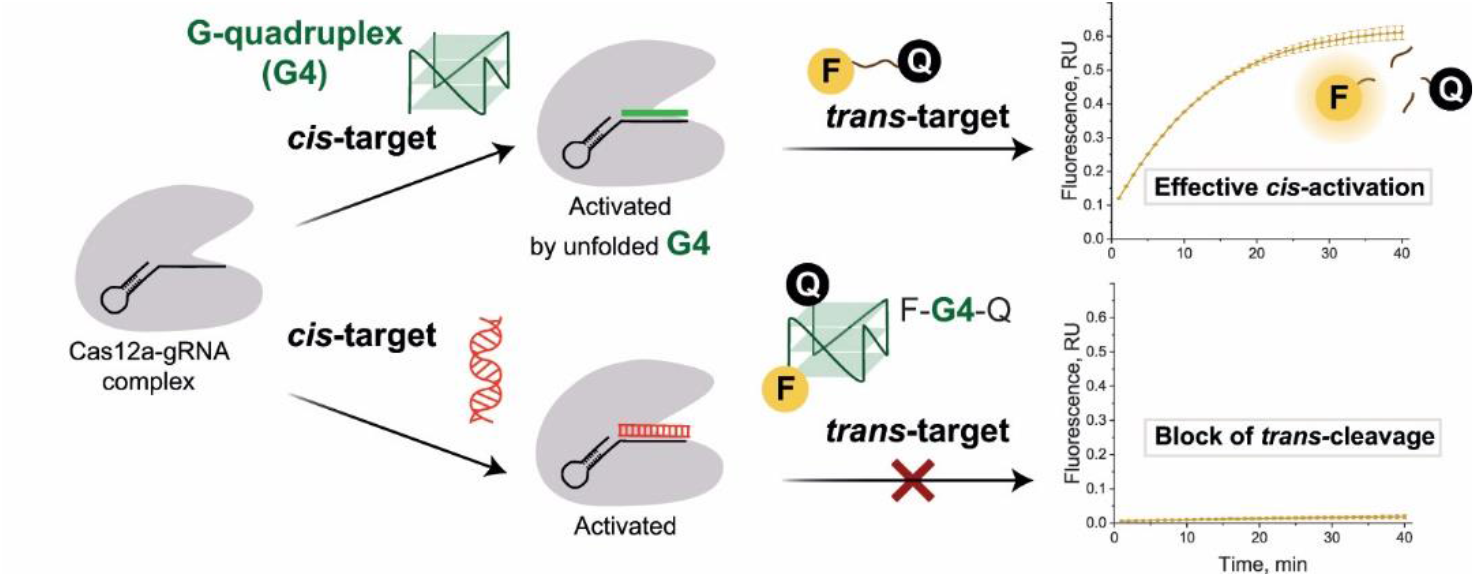

## Introduction

Cas12a is an RNA-guided CRISPR nuclease widely used in nucleic-acid diagnostics because recognition of a sequence-matched *cis*-DNA-target activates nonspecific *trans*-cleavage of single-stranded DNA (ssDNA) [1,2]. The *cis*-target is the nucleic acid recognized by the guide RNA–Cas12a complex; independent ssDNA molecules cleaved after activation are *tran*s-targets, and labelled *trans*-targets serve as reporters. Because reporter cleavage generates the diagnostic signal, changes in reporter sequence, length or folding can directly alter Cas12a *trans*-cleavage efficiency and signal output [3–5].

Canonical Cas12a activation proceeds through a PAM-containing double-stranded DNA (dsDNA) target, but complementary ssDNA can also activate Cas12a in a PAM-independent manner [6,7]. For such ssDNA *cis*-targets, intramolecular folding can be more consequential than for canonical duplex targets: a folded *cis*-target must unfold sufficiently to form the guide-target hybrid required for Cas12a activation [8,9]. Folding can therefore affect both signal-generating steps by limiting guide-target hybrid formation in *cis*-targets or by changing *trans*-target accessibility to collateral cleavage.

DNA targets with defined secondary structure are therefore of interest as potential regulators of CRISPR/Cas12a systems. G-quadruplexes (G4s) represent one such highly ordered class of DNA structures with condition-dependent stability. G4s are non-canonical four-stranded structures formed by guanine-rich sequences; stacked G-tetrads are held together by Hoogsteen hydrogen bonding and stabilized by coordinated monovalent cations, most effectively K^+^ [10]. Their folding and stability depend on sequence, loop architecture, ionic composition and temperature, making G4s controllable structural elements. This controllability contributes to their regulatory roles in transcription, replication and genome stability [11,12] and has been exploited in engineered DNA systems, including aptamers [13], catalytic DNAzymes [14] and G4-based biosensing reporters [4,5,15].

In Cas12a systems, G4 motifs have so far been examined mainly as *trans*-targets or reporter-derived signal elements. Activated Cas12a cleaves telomeric and thrombin-binding G4 *trans*-targets, although G4 stability can markedly prolong cleavage [16]. Later reporter designs used G4 stability, potassium responsiveness and topology to modulate collateral-cleavage signals, including pre-folded [4], molecular-beacon [5], phosphorothioate-modified [17], split [18] and G-triplex [19] formats. These studies showed that G4 folding can slow or tune Cas12a *trans*-cleavage and can suppress it under high-salt reporter conditions, but they did not establish whether a compact parallel G4 scaffold can fully resist collateral cleavage. It also remained unknown how a folded G4 ssDNA affects Cas12a when used as a guide-complementary *cis*-DNA-target.

Evidence for interactions between CRISPR nucleases or nuclease-dead variants and G4-containing DNA is available primarily from Cas9-based systems. Targeting promoter G4 motifs with CRISPR–dCas9 can stabilize or destabilize folding in a strand-orientation-dependent manner and modulate transcription [20,21]. Other studies reported Cas9/dCas9 interactions with folded G4 targets [22,23] and selective targeting of individual G4 and i-motif structures by chemically modified Cas9 [24]. Together, these studies show that guide-programmed CRISPR complexes can recognize G4-containing DNA, but they do not establish whether a folded ssDNA G4 can serve as a Cas12a *cis*-DNA-target.

Accordingly, the influence of G4 folding on each Cas12a functional mode remains to be defined. In the *cis* context, G4-containing *cis*-targets reveal whether guide-directed recognition can unfold a compact structure and how fold stability changes Cas12a activation. In the *trans* context, G4-containing *trans*-targets reveal whether the folded backbone remains accessible to the activated nonspecific nuclease and how strongly folding suppresses collateral cleavage.

To address this gap, compact quadruplexes based on contiguous (TGGG)n repeats were selected as a defined model system. Closely related d(GGGT)_4_-family aptamers, historically described as T30923/T30695, bind HIV-1 integrase and the interleukin-6 receptor and are structurally well characterized as parallel G4s with three G-tetrads and single-thymidine reversed-chain loops [25–28]. Because (TGGG)n repeats share this compact parallel architecture, they provide a matched scaffold for comparing guide-complementary *cis*-targets with independent labelled *trans*-target reporters.

Here, we examined compact (TGGG)n G4s as *cis*-DNA-targets and labelled *trans*-target reporters. The same architecture was tested with LbCas12a and AsCas12a using real-time cleavage assays, circular dichroism (CD), FRET and denaturing gel electrophoresis. The data show a strong functional asymmetry.

Compact parallel G4s are competent *cis*-targets that activate Cas12a after guide-dependent unfolding, but the same folded motifs resist collateral *trans*-cleavage. Disrupting the core restores *trans*-cleavage, whereas K^+^-mediated stabilization suppresses *cis*-activation. Antiparallel G4s show partial, rather than complete, *trans*-protection. These results define G4 folding as a tunable structural gate that differentially controls Cas12a activation and collateral cleavage.

## Materials and Methods

### Materials

DNA oligonucleotides, including unmodified G-quadruplex-forming sequences, fluorophore/quencher-labelled reporters, FRET constructs, complementary strands and other oligonucleotides (the sequences are presented in Supplementary Information (SI), Table S1), were synthesized by Lumiprobe (Moscow, Russia) and Evrogen (Moscow, Russia). Guide RNAs (SI, Table S1) were synthesized by Syntol (Moscow, Russia). LbCas12a, Q5 polymerase (2 U/µL), Exonuclease I (20 U/µL), DNase I (2 U/µL) and the supplied 10× buffers were from New England Biolabs (Ipswich, MA, USA). AsCas12a was from GenScript (Nanjing, China). DNA electrophoresis markers, dNTPs and the Cleanup Standard DNA extraction kit were purchased from Evrogen (Moscow, Russia). Agarose was from neoFroxx (Einhausen, Germany), and Amicon Ultra 3K filters were from Merck Millipore (Burlington, MA, USA). All salts and organic compounds were of analytical grade.

### General reaction conditions

Unless otherwise indicated, Cas12a reactions were performed in NEBuffer 2.1 (New England Biolabs; 50 mM NaCl, 10 mM Tris-HCl, 10 mM MgCl_2_, 100 µg/mL BSA, pH 7.9). G4 folding and protein-free CD and fluorescence/FRET measurements were performed in the same buffer without BSA, referred to here as BSA-free NEBuffer 2.1. Where indicated, these buffers were supplemented with KCl or LiCl (10 or 50 mM final) or prepared without MgCl_2_ or BSA, as specified.

To prepare folded G4 samples, G-quadruplex-forming oligonucleotides were heated at 95 °C for 10 min in BSA-free NEBuffer 2.1, with the indicated salt modifications, and then cooled at room temperature for 15 min to allow G4 formation.

### Preparation of the double-stranded DNA *cis*-target (activator) for CRISPR–Cas12a reaction

A 503-bp fragment of the 16S-23S intergenic spacer region of *Clavibacter sepedonicus* was used as a standard double-stranded DNA *cis*-activator for CRISPR–Cas12a reactions. The guide RNA was designed with CHOPCHOP (https://chopchop.cbu.uib.no/), and PCR primers (SI, Table S1 and Figure S1) were designed with Primer-BLAST (https://www.ncbi.nlm.nih.gov/tools/primer-blast/).

Cells of *C. sepedonicus* were obtained from the bacterial collection of the All-Russian Plant Quarantine Center (Bykovo, Moscow Region, Russia). Total DNA was isolated using the PhytoSorb kit (Syntol, Moscow, Russia). DNA concentration was measured spectrophotometrically using a NanoDrop ND-2000 instrument (Thermo Fisher Scientific, USA).

Preparative PCR (38 cycles) from total genomic DNA was carried out in a Bio-Rad T100 thermal cycler (Bio-Rad, USA) at 95 °C for 30 s (denaturation), 55 °C for 30 s (annealing) and 72 °C for 60 s (extension) using primers PCRCs-F/PCRCs-R. The final PCR mixture contained 500 nM of each primer, 200 µM dNTPs, Q5 polymerase (2 U) and 20 ng of total DNA. Amplicons were separated in a 2% agarose gel (20 mM Tris-acetate buffer, 0.2 mM EDTA, pH 8.3) with ethidium bromide. The gel was imaged under UV illumination using a Clinx gel documentation system (Clinx Science Instruments, China) (SI, Figure S2), and gel slices corresponding to the 503-bp product were excised. dsDNA was purified from the gel using the Evrogen kit, and its concentration was measured with a NanoDrop ND-2000 instrument.

### CRISPR–Cas12a assays

Cas12a collateral-cleavage activity was monitored in real time by fluorescence dequenching of ssDNA reporters carrying fluorophore and quencher labels, hereafter fluorophore/quencher called *trans*-targets. The Cas12a–gRNA complex was assembled by pre-incubating Cas12a with the corresponding gRNA in 1× NEBuffer 2.1 for 10 min at 25 °C. The *trans*-target (500 nM final) and the *cis*-activator (dsDNA amplicon or single-stranded *cis*-target with final concentrations of 60 nM Cas12a, 60 nM gRNA and 500 nM *trans*-target. Fluorescence was recorded for 40 min at 37 or 47 °C on a LightCycler 96 system (Roche, Switzerland) using the ROX/Texas Red or FAM channel. Water was used instead of the activator in no-target controls. R(T)_15_, an ssDNA reporter containing 15 thymidine residues and ROX/BHQ2 labels, was used as the standard reference reporter for LbCas12a; R-FAM/BHQ1 was used for AsCas12a (SI, Table S1). When G-quadruplexes, their mutants and G-rich oligonucleotides were tested as *trans*-targets, the 503-bp dsDNA fragment was used as the *cis*-target. When G-quadruplexes and their mutants were tested as *cis*-targets, R(T)_15_ or R-FAM/BHQ1 was used as the *trans*-target.

Specificity of guide-dependent *cis*-activation was assessed using the core-disrupting mutant (TGGG)_5_-M1, as well as the fold-preserving long-loop mutant (TGGG)_4_-M2 and shortened (TGGG)_2_/(TGGG)_3_ oligonucleotides (SI, Table S1). For concentration-dependence experiments, the *cis*-target was serially titrated and fluorescence intensity at 40 min was plotted against target concentration.

### Exonuclease I and DNase I cleavage assays for G4 and linear reporters

Reporter susceptibility to non-CRISPR nucleases was tested by incubating 500 nM reporter with Exonuclease I (40 U/reaction) or DNase I (4 U/reaction) in the corresponding 1× buffer at 37 °C, with real-time fluorescence detection on a LightCycler 96 system (Roche, Switzerland). For DNase I experiments, G-quadruplex reporters were assayed either alone or after hybridization with complementary strands. Reporter and complementary strand (1:2) were annealed by heating to 95 °C for 10 min and cooling at room temperature for 15 min.

Calibration standards for R(T)_15_ and R-FAM/BHQ1 were prepared by complete Exonuclease I digestion of 2 µM reporter for 80 min at 37 °C in the corresponding 1× buffer. Reactions were stopped with EDTA (5 mM final). The digested reporters were diluted to 7.8 nM-1 µM and used to generate calibration curves relating fluorescence intensity to cleaved reporter concentration. The fluorescence curves and calibration curves are presented in SI, Figure S3.

### Determination of CRISPR–Cas12a kinetic parameters for *trans*-cleavage

Initial reaction velocity (V_0_) was used to compare Cas12a activation and reporter cleavage under different conditions. V_0_ was calculated as the slope of the linear region of kinetic fluorescence curves recorded for 40 min at 37 or 47 °C on a LightCycler 96 system using the ROX/Texas Red or FAM channel. Fluorescence intensity was first converted to cleaved reporter concentration using the Exonuclease I calibration curves (SI, Figure S3).

Kinetic parameters of *trans*-cleavage by Cas12a activated with the (TGGG)_5_ *cis*-target were determined using a modified Ramachandran and Santiago protocol [2]. The gRNA–Cas12a complex (5.5 nM) was activated with 42 nM (TGGG)_5_ *cis*-target for 30 min at 37 °C. The reporter (R(T)_15_ for LbCas12a or R-FAM/BHQ1 for AsCas12a) was then titrated from 62.5 to 3000 nM. V_0_ values were plotted against reporter concentration and fitted with the Michaelis–Menten function to estimate K_M_ (Michaelis–Menten constant), k_cat_ (enzyme turnover rate) and the catalytic efficiency k_cat_/K_M_.

### Circular dichroism spectroscopy and thermal denaturation of G4

CD spectra were recorded on a Chirascan spectrometer (Applied Photophysics, Leatherhead, Surrey, UK) using a 5-mm path-length cuvette, a 1.5-nm bandwidth and a 3-s integration time. Oligonucleotides (1 µM) were measured in BSA-free NEBuffer 2.1, with or without KCl (10 or 50 mM) or LiCl (50 mM), at 20, 37, 47 and 85 °C over 210–340 nm. UV absorbance spectra were recorded using the same instrument. CD spectra were converted from ellipticity (mdeg) to mean residue molar ellipticity, [θ]_MR_ (deg cm^2^ dmol^−1^), using [θ] = 100 × θ/(c × L), followed by normalization to nucleotide number. Thermal denaturation was followed at 260 nm from 20 to 77 °C in BSA-free NEBuffer 2.1, with or without 50 mM KCl or 50 mM LiCl.

### FRET measurements of G4 as a *cis*-activator

Doubly end-labelled (TGGG)_5_ constructs bearing donor/acceptor pairs AF555-(TGGG)_5_-AF647, FAM-(TGGG)_5_-TAMRA, Coumarin343-(TGGG)_5_-FAM, Cy3-(TGGG)_5_-Cy5 and JOE-(TGGG)_5_-Cy5, as well as the single-fluorophore/dark-quencher construct BHQ-(TGGG)_5_-Cy5 (SI, Table S1), were folded as described above in BSA-free NEBuffer 2.1. The starting concentration of the labelled construct was 1 µM. 2D emission spectra and 3D excitation-emission maps were acquired on a Shimadzu RF-6000 spectrofluorophotometer (Kyoto, Japan). For 3D maps, excitation/emission ranges were 450–750/500–750 nm for all pairs except Coumarin343-(TGGG)_5_-FAM, for which the ranges were 350–650/400–650 nm. 2D spectra were recorded across the emission range at the donor-excitation maximum identified from the corresponding 3D map.

For each donor/acceptor construct, 3D maps were used to identify the donor- and acceptor-excitation maxima and the corresponding emission maxima. Fluorescence intensity (FI) values were then extracted from 2D spectra recorded at the donor-excitation maximum. FI(FRET)/FI(donor) was calculated from the FI values at these emission maxima: the sensitized acceptor-emission maximum divided by the donor-emission maximum in the same donor-excited spectrum. Folded constructs were then compared with duplex reference samples prepared by adding complementary (CCCA)_5_ at a 1:2 G4:complement ratio. For JOE-(TGGG)_5_-Cy5, duplex formation was further characterized by titrating (CCCA)_5_ at G4:complement ratios of 1:0.5, 1:0.75, 1:1, 1:2 and 1:3. The folded construct and each titration point were prepared with 1 µM labelled construct before dilution and recorded as undiluted and 2-, 3- and 5-fold diluted samples. These dilution series confirmed that FI(FRET)/FI(donor) was independent of total fluorescence intensity within the measured range. Constructs with the largest separation between folded and duplex states were selected for CRISPR–Cas12a assays.

After selection of JOE-(TGGG)_5_-Cy5 as the main FRET construct, additional 3D maps were recorded for its complexes with gRNA-20 alone and with gRNA-20/AsCas12a. The construct:gRNA and construct:gRNA:AsCas12a mixtures were incubated for 60 min before 3D map acquisition. In both series, all components were used at equimolar concentrations of 200, 250 or 400 nM. FI(FRET)/FI(donor) was calculated from these 3D maps at the donor-excitation maximum using the emission maxima defined above. For FRET datasets reported as mean ± SD, values were calculated from the corresponding dilution or concentration series because FI(FRET)/FI(donor) did not depend on dilution.

After spectral measurements, R-FAM/BHQ1 reporter (1 µM) was added to confirm AsCas12a trans-nuclease activation by kinetic fluorescence recording.

### Estimation of G4 as a *cis*-activator using polyacrylamide gel electrophoresis

Reaction mixtures containing FAM-(TGGG)_5_ *cis*-target (300 nM), R-FAM/BHQ1 reporter (1 µM) and gRNA/Cas12a at 60/60 or 300/300 nM were analyzed by vertical PAGE in a Mini-PROTEAN system (Bio-Rad Laboratories, Hercules, CA, USA). Electrophoresis was performed in a 15% polyacrylamide gel (acrylamide:methylenebisacrylamide, 29:1) under denaturing conditions (40 mM Tris-acetate buffer, pH 8.3, 1 mM EDTA, 7 M urea). Samples were mixed with an equal volume of 40% glycerol/0.01% bromophenol blue/0.01% xylene cyanol in 40 mM Tris-acetate buffer with 0.4 mM EDTA, pH 8.3. Samples were loaded directly onto the gel and run for 2 h at a constant power of 1 W. Gels were imaged by in-gel FAM fluorescence using a ChemiDoc XRS+ system (Bio-Rad, USA), and images were processed using Image Lab 6.0.1 software (Bio-Rad Laboratories).

### Statistics and data analysis

Unless otherwise stated, experiments were performed with n = 3 replicates and data are presented as mean ± standard deviation (SD). Data were processed and visualized by OriginPro 2021 (OriginLab, Northampton, MA, USA).

## Results

### A compact (TGGG)n quadruplex core activates Cas12a as *cis*-target but resists collateral *trans*-cleavage

Compact (TGGG)n motifs fold into intramolecular parallel quadruplexes in which four G-tracts stack into three G-tetrads [25,26]. The experimental set included (TGGG)_4_, the extended (TGGG)_5_, the TATA-tailed (TGGG)_4_-TATA, and the core-disrupting mutant (TGGG)_5_-M1 (SI, Table S1), whose substitutions of G to A nucleotides abolish the quadruplex. These related constructs were analyzed in two positions of the Cas12a reaction: as labelled *trans*-target reporters for collateral *trans*-cleavage and as guide-complementary *cis*-targets for nuclease activation. Both activities were examined with LbCas12a and AsCas12a.

First, (TGGG)n oligonucleotides were tested as labelled *trans*-target reporters as shown schematically in Fig. 1A. In this assay, Cas12a was activated by the 503-bp *C. sepedonicus* 16S-23S spacer fragment used as the *cis*-dsDNA-target (activator), together with its cognate gRNA. The G4 reporters carried ROX and BHQ labels at opposite ends (Fig. 1B).

**Figure 1.**
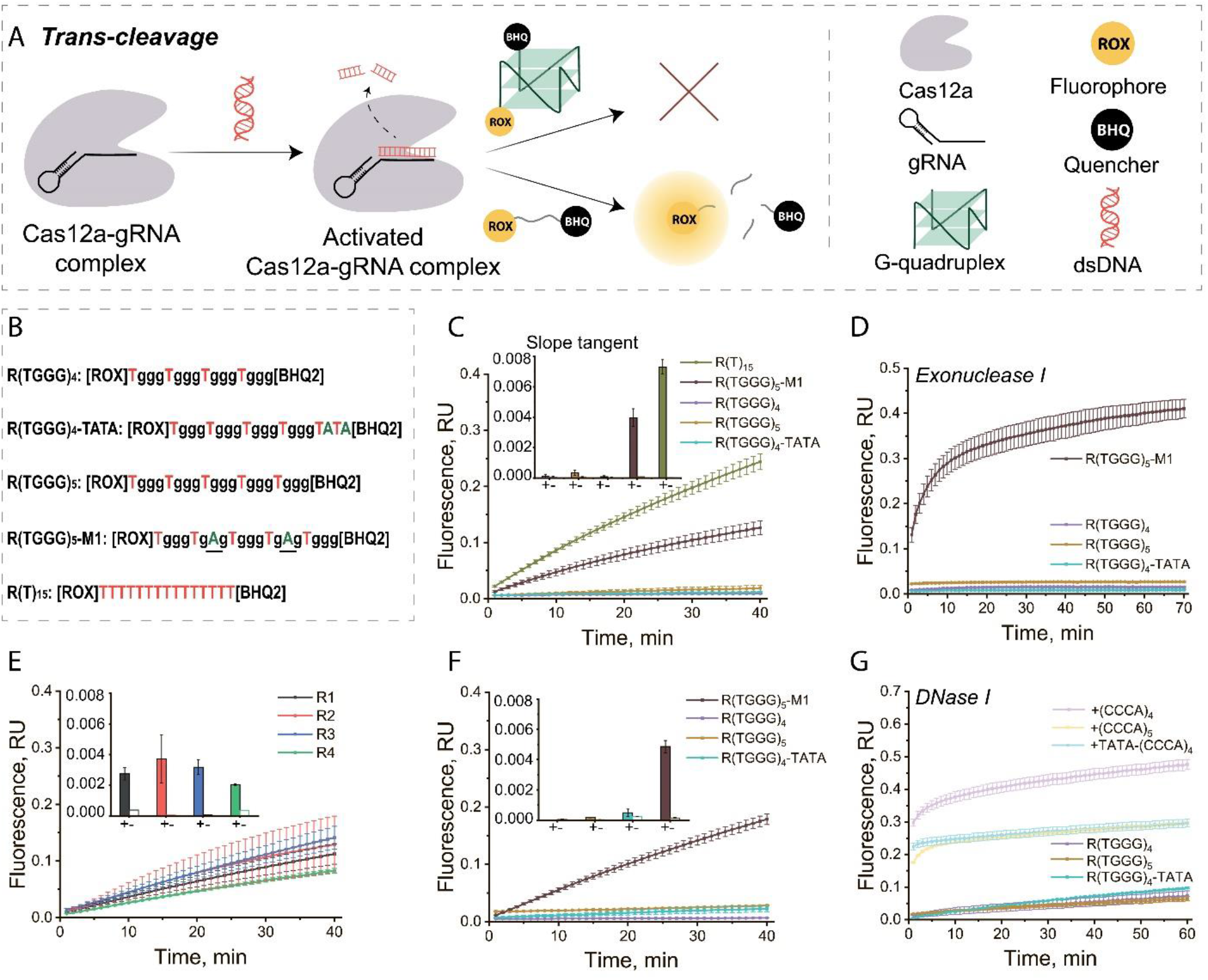
Compact parallel G4 reporters resist collateral *trans*-cleavage. (A) Scheme of the CRISPR– Cas12a reaction with a G-quadruplex reporter as the *trans*-target. (B) Sequences of R(TGGG)_4_, R(TGGG)_4_-TATA, R(TGGG)_5_, R(TGGG)_5_-M1 and the standard R(T)_15_ reporter. (C) Fluorescence curves for R(TGGG)_4_, R(TGGG)_5_, R(TGGG)_4_-TATA and R(TGGG)_5_-M1 compared with R(T)_15_ (LbCas12a, 37 °C); inset, slope of the initial linear phase for activated reactions and no-activator controls. (D) Cleavage of R(TGGG)_4_, R(TGGG)_5_, R(TGGG)_4_-TATA and R(TGGG)_5_-M1 by Exonuclease I at 37 °C. (E) Fluorescence curves for G-rich non-G4 reporters R1–R4 (LbCas12a, 37 °C); inset, initial slope. (F) Fluorescence curves for R(TGGG)_4_, R(TGGG)_5_, R(TGGG)_4_-TATA and R(TGGG)_5_-M1 for AsCas12a at 37 °C. (G) DNase I cleavage of R(TGGG)_4_, R(TGGG)_5_ and R(TGGG)_4_-TATA before and after hybridization with complementary strands at 37 °C.

At 37 °C, the quadruplex reporters R(TGGG)_4_, R(TGGG)_5_ and R(TGGG)_4_-TATA did not increase fluorescence above the no-activator control, whereas the linear reference R(T)_15_ and the unfolded mutant R(TGGG)_5_-M1 were rapidly cleaved by activated LbCas12a (Fig. 1C). G-rich but non-quadruplex controls R1–R4 (SI, Table S1) were also efficiently cleaved (Fig. 1E), showing that resistance was associated with quadruplex folding rather than guanine richness alone. The same pattern was observed with AsCas12a (Fig. 1F) and at 47 °C (SI, Figure S4), indicating that the effect is robust across the working temperature range of Cas12a.

Additional nuclease-accessibility controls were used to examine whether the absence of reporter signal reflected a labelling artifact or reduced access to the folded G4. Exonuclease I, a processive 3′→5′ exonuclease that initiates from an accessible single-stranded 3′ end [29], did not measurably cleave folded R(TGGG)n reporters but almost completely degraded the unfolded mutant R(TGGG)_5_-M1 (Fig. 1D). DNase I produced only a weak signal with pre-folded quadruplex reporters. In contrast, after hybridization with complementary strands ((CCCA)_4_, (CCCA)_5_ or TATA-(CCCA)_4_), which converted the G-rich strand into a duplex, DNase I cleavage produced a steep fluorescence increase (Fig. 1G). These controls indicate that the labels were functional and that limited backbone accessibility accounts for the resistance of folded (TGGG)n reporters to the cleavage.

Second, the same (TGGG)n oligonucleotides were examined as guide-complementary *cis*-targets (scheme, Fig. 2A; gRNA/target pairs, Fig. 2B). All three folded structures activated LbCas12a (R(T)_15_ readout; Fig. 2C) and AsCas12a (R-FAM/BHQ1 readout; Fig. 2D), causing cleavage of standard reporters. Activation was guide-specific: (TGGG)_5_-M1 activated LbCas12a with the cognate gRNA-20M1 but not with non-cognate gRNA-20 (Fig. 2C). At 47 °C, LbCas12a retained similar activity toward the G4 *cis*-targets, whereas AsCas12a activity was reduced under these conditions (SI, Figure S5). Initial velocities (V_0_) showed that activation was lower at 47 °C than at 37 °C for both orthologs and that (TGGG)_5_ activated the reaction more slowly than (TGGG)_4_ or (TGGG)_4_-TATA (Fig. 2E). Because (TGGG)_5_ was the slowest compact *cis*-DNA-target, this scaffold was used for Michaelis–Menten analysis at 37 °C. The (TGGG)_5_-activated reactions gave similar k_cat_ values for LbCas12a and AsCas12a (2.9 and 2.79 min^−1^, respectively), with k_cat_/K_M_ values of 5.9×10^4^ and 8.9×10^4^ M^−1^ s^−1^. The higher apparent catalytic efficiency of AsCas12a resulted from its lower K_M_ (0.52 µM versus 0.82 µM for LbCas12a) (SI, Figure S6).

**Figure 2.**
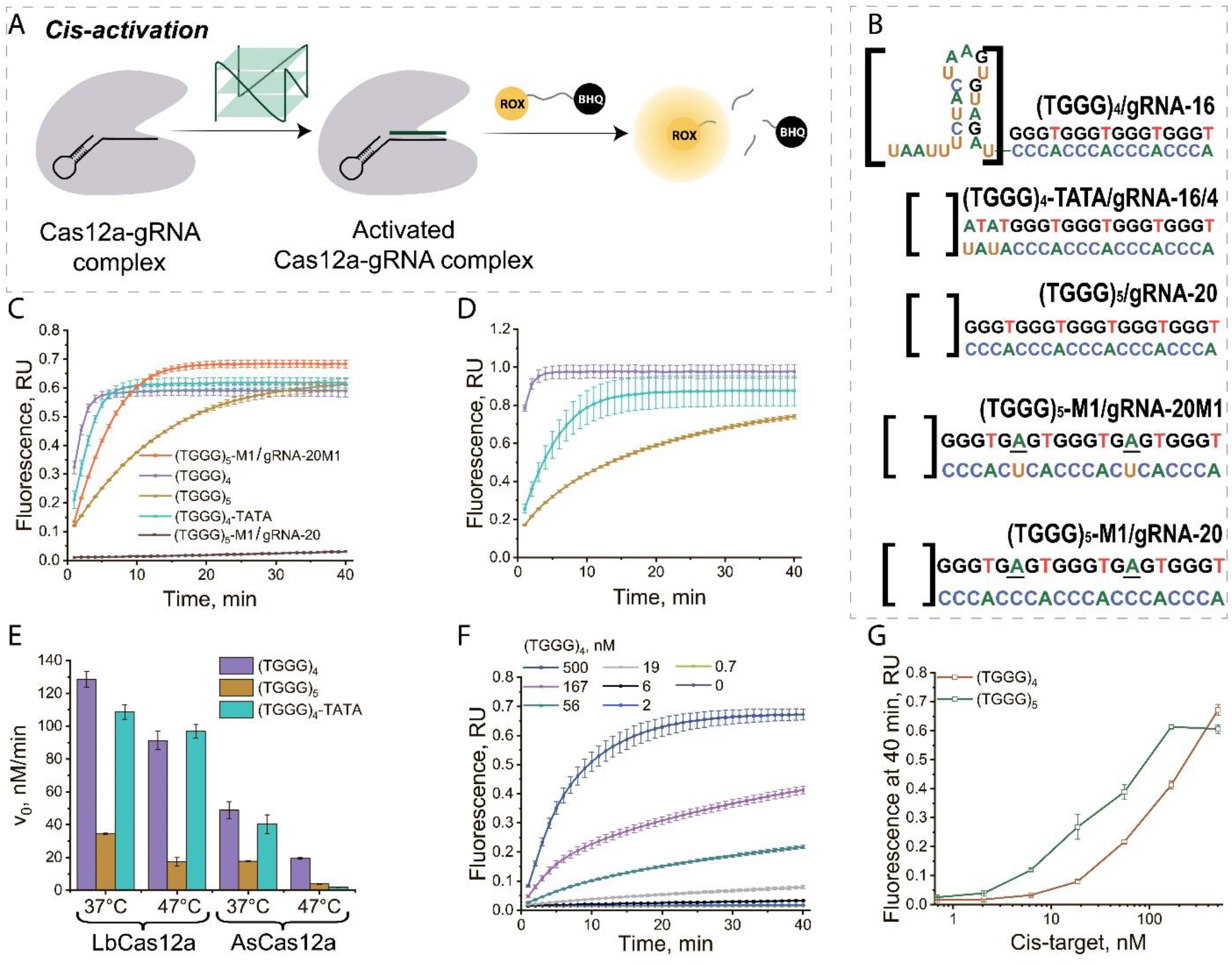
Compact parallel G4s activate Cas12a as *cis*-targets. (A) Scheme of the CRISPR–Cas12a reaction with a G-quadruplex as the *cis*-target. (B) Cognate gRNA/*cis*-target pairs: (TGGG)_4_/gRNA-16, (TGGG)_4_-TATA/gRNA-16/4, (TGGG)_5_/gRNA-20, (TGGG)_5_-M1/gRNA-20M1 and (TGGG)_5_-M1/gRNA-20. (C) Fluorescence curves for (TGGG)_4_, (TGGG)_5_, (TGGG)_4_-TATA and (TGGG)_5_-M1 with gRNA-20M1 or gRNA-20 (LbCas12a, 37 °C, R(T)_15_ readout). (D) As in (C), for AsCas12a (37 °C, R-FAM/BHQ1 readout). (E) Initial velocities V_0_ for (TGGG)_4_, (TGGG)_5_ and (TGGG)_4_-TATA with LbCas12a and AsCas12a at 37 and 47 °C. (F) Fluorescence curves (LbCas12a, R(T)_15_ readout) for different concentrations of (TGGG)_4_ with gRNA-16 at 37 °C. (G) Fluorescence intensity at 40 min versus *cis*-target concentration for (TGGG)_4_ and (TGGG)_5_. Data are mean ± SD (n = 3).

The sequence-length requirement for *cis*-activation was evaluated with shortened non-quadruplex oligonucleotides, (TGGG)_2_ and (TGGG)_3_. The strongest contrast was observed with gRNA-16/4: the short oligonucleotides produced little activation (SI, Figure S7). Across the tested gRNAs, (TGGG)_2_ did not activate Cas12a, whereas the 12-nt (TGGG)_3_ oligonucleotide produced detectable activation with gRNA-16 and gRNA-20. At 47 °C, the full-length G4-forming oligonucleotides gave higher signals than (TGGG)_3_. This length dependence is consistent with a requirement for productive gRNA–*cis*-target pairing initiated through a short seed segment [8,9], rather than with nonspecific activation by repeated TGGG motifs.

*Cis*-activation also depended on the amount of G4 target: lowering the concentration of (TGGG)_4_ or (TGGG)_5_ reduced reporter cleavage (Fig. 2F and SI, Figure S8). Endpoint concentration curves built from 40-min fluorescence values allowed detection of (TGGG)_5_ down to 2 nM and (TGGG)_4_ down to 6 nM (Fig. 2G).

Together, these data establish a functional asymmetry: compact (TGGG)n G4s resist collateral cleavage as *trans*-target reporters but activate Cas12a as guide-complementary *cis*-targets. The effect is not explained by G-rich sequence alone, because G-rich non-G4 controls are cleaved and disruption of the core restores trans-cleavage. The following sections present four analyses motivated by this finding:

1. characterization of (TGGG)n topology and stability under CRISPR–Cas12a conditions by CD;
2. testing of fold-stability effects on *cis*-activation by varying the ionic environment (K^+^/Li^+^);
3. direct monitoring of conformational changes and cleavage of the G4 *cis*-target;
4. 4 and evaluation of whether the asymmetry is retained in antiparallel G4s.

### (TGGG)n fold into parallel G-quadruplexes whose stability is tunable by ions and temperature

The functional asymmetry observed for (TGGG)n required structural characterization under buffer conditions matching the Cas12a assays. At 37 °C in BSA-free NEBuffer 2.1, all three oligonucleotides produced CD spectra with a positive band at 262–264 nm and a negative band near 240 nm (Fig. 3A), consistent with parallel G4 topology. The schematic models in Fig. 3B therefore represent parallel three-G-tetrad folds consistent with the CD signatures and with established structures of related d(GGGT)_4_-family quadruplexes [25,26]. The additional TGGG repeat in (TGGG)_5_ and the terminal TATA segment in (TGGG)_4_-TATA preserved the parallel CD signature, but changed the band amplitude. The positive CD band was strongest for (TGGG)_5_ and weakest for (TGGG)_4_-TATA, suggesting differences in the population and/or ordering of the parallel G4 state under the same conditions. This trend is consistent with the activation series in Fig. 2E: the oligonucleotide with the strongest parallel G4 signature, (TGGG)_5_, showed the lowest initial rates, as expected if *cis*-activation requires disruption of the folded G4 and formation of the gRNA–target complex.

**Figure 3.**
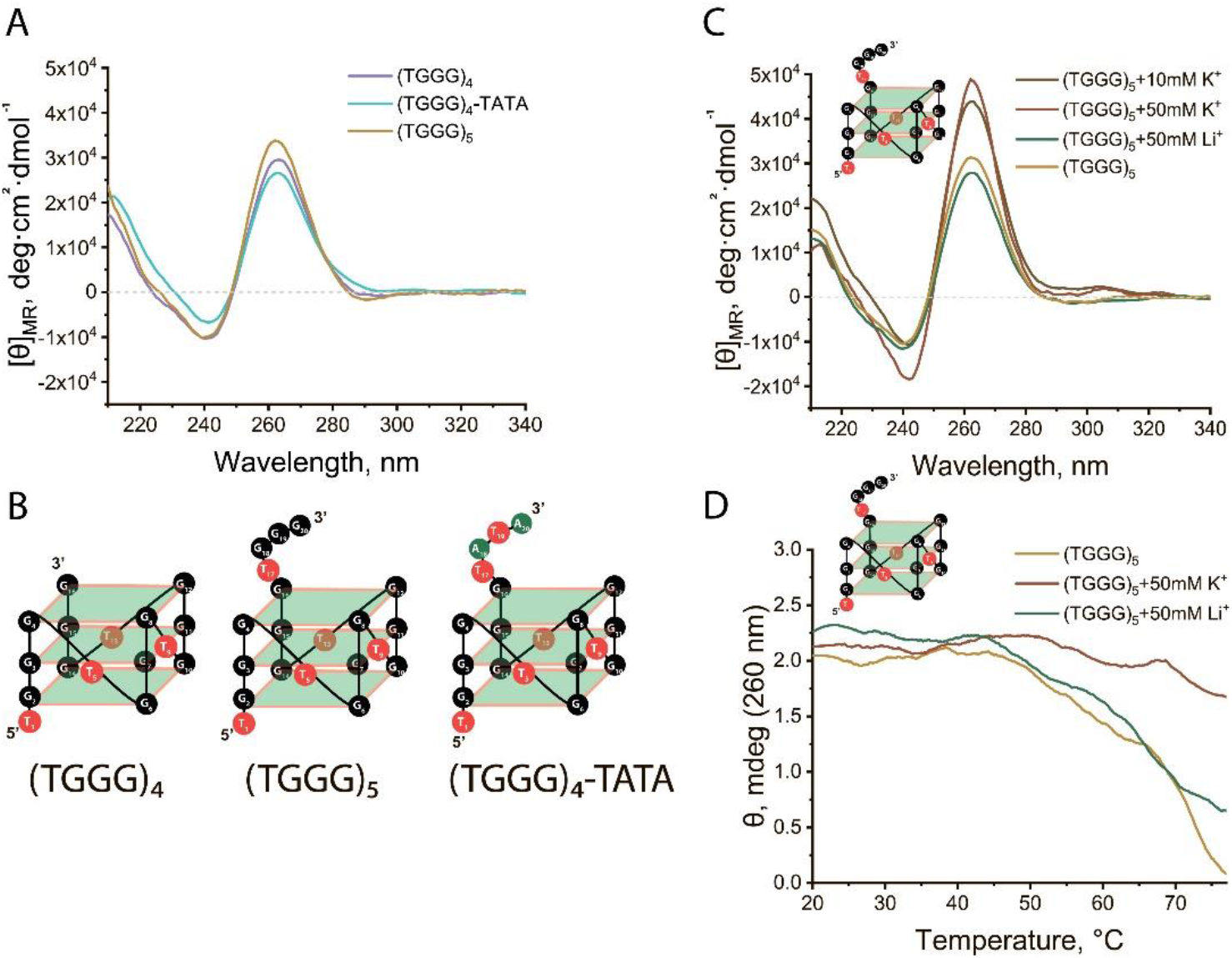
(TGGG)n oligonucleotides form parallel G-quadruplexes with ion- and temperature-dependent stability. (A) CD spectra of (TGGG)_4_, (TGGG)_5_ and (TGGG)_4_-TATA at 37 °C in NEB buffer. (B) Structural models of the parallel quadruplexes based on three stacked G-tetrads in related d(GGGT)_4_-family structures. (C) CD spectra of (TGGG)_5_ in NEB and with 10 mM KCl, 50 mM KCl or 50 mM LiCl at 37 °C. (D) Melting curves of (TGGG)_5_ at 260 nm in NEB and with 50 mM KCl or 50 mM LiCl.

Because monovalent-cation coordination provides a controlled way to vary G4 stability, cation effects were examined for the compact (TGGG)_5_ scaffold in Cas12a-compatible buffer (Fig. 3C). KCl increased the positive 263-nm band in a concentration-dependent manner between 10 and 50 mM, consistent with K^+^ coordination between G-tetrad planes and stabilization of the fold. LiCl (50 mM) produced a weaker parallel G4 signature than KCl, consistent with weaker G-tetrad stabilization by Li^+^. A similar increase in the 263-nm band upon addition of 10 mM KCl was observed for (TGGG)_4_ and (TGGG)_4_-TATA (SI, Figure S9D–G).

Thermal denaturation monitored at 260 nm over 20–77 °C further supported ion-dependent stabilization of (TGGG)_5_ (Fig. 3D). In BSA-free NEBuffer 2.1, the signal began to decrease above ∼50 °C and approached baseline by the end of the temperature ramp, consistent with substantial loss of the parallel fold. LiCl (50 mM) produced an intermediate profile: the signal decreased over a similar temperature range but remained higher than in BSA-free NEBuffer 2.1 at 77 °C. In 50 mM KCl, the 260-nm signal was largely retained across the measured range and decreased only modestly at the highest temperatures. CD spectra recorded at 85 °C were consistent with this stability order: the 263-nm band remained prominent in 50 mM KCl but was strongly reduced in BSA-free NEBuffer 2.1 and in 50 mM LiCl (SI, Figure S9C). Temperature series at 20, 37 and 47 °C showed lower 263-nm band amplitudes upon heating in BSA-free NEBuffer 2.1, whereas 10 mM KCl largely preserved the parallel G4 signature for (TGGG)_5_, (TGGG)_4_ and (TGGG)_4_-TATA (SI, Figure S9). Thus, K^+^ provided the strongest thermal stabilization of the compact parallel fold under the assay conditions.

Together, the CD and melting data place the functional asymmetry observed above in a defined structural context: the tested (TGGG)n sequences form compact parallel G4s under Cas12a-compatible conditions. Their stability is strongly enhanced by K^+^, partially supported by Li^+^ and weakened by heating. This ion- and temperature-dependent structural series provided the basis for testing whether stabilization of the G4 *cis*-target limits Cas12a activation.

### Control of Cas12a *cis*-activation through G-quadruplex folding stability

The CD and melting experiments showed that (TGGG)n G4 stability was sensitive to monovalent ions and temperature. The next experiments therefore tested Cas12a activation under conditions that either stabilized or weakened the folded *cis*-target. If stabilization limits G4 unfolding, formation of the gRNA–target hybrid should become less efficient, leading to reduced Cas12a activation. As a control for salt effects on the Cas12a and standard reporter cleavage, the 503-bp *cis*-dsDNA-target (activator) was tested with the R(T)_15_ *trans*-target reporter under the same ionic conditions. At 37 °C, addition of 50 mM KCl or 50 mM LiCl did not substantially suppress dsDNA-dependent activation, and omission of MgCl_2_ abolished activity (Fig. 4A). At 47 °C, KCl slightly increased the dsDNA-dependent signal, LiCl produced only a modest decrease, and omission of MgCl_2_ again abolished activity (SI, Figure S10). Thus, the monovalent salts used to modulate G4 stability did not inhibit LbCas12a activation by a canonical *cis*-dsDNA-target.

**Figure 4.**
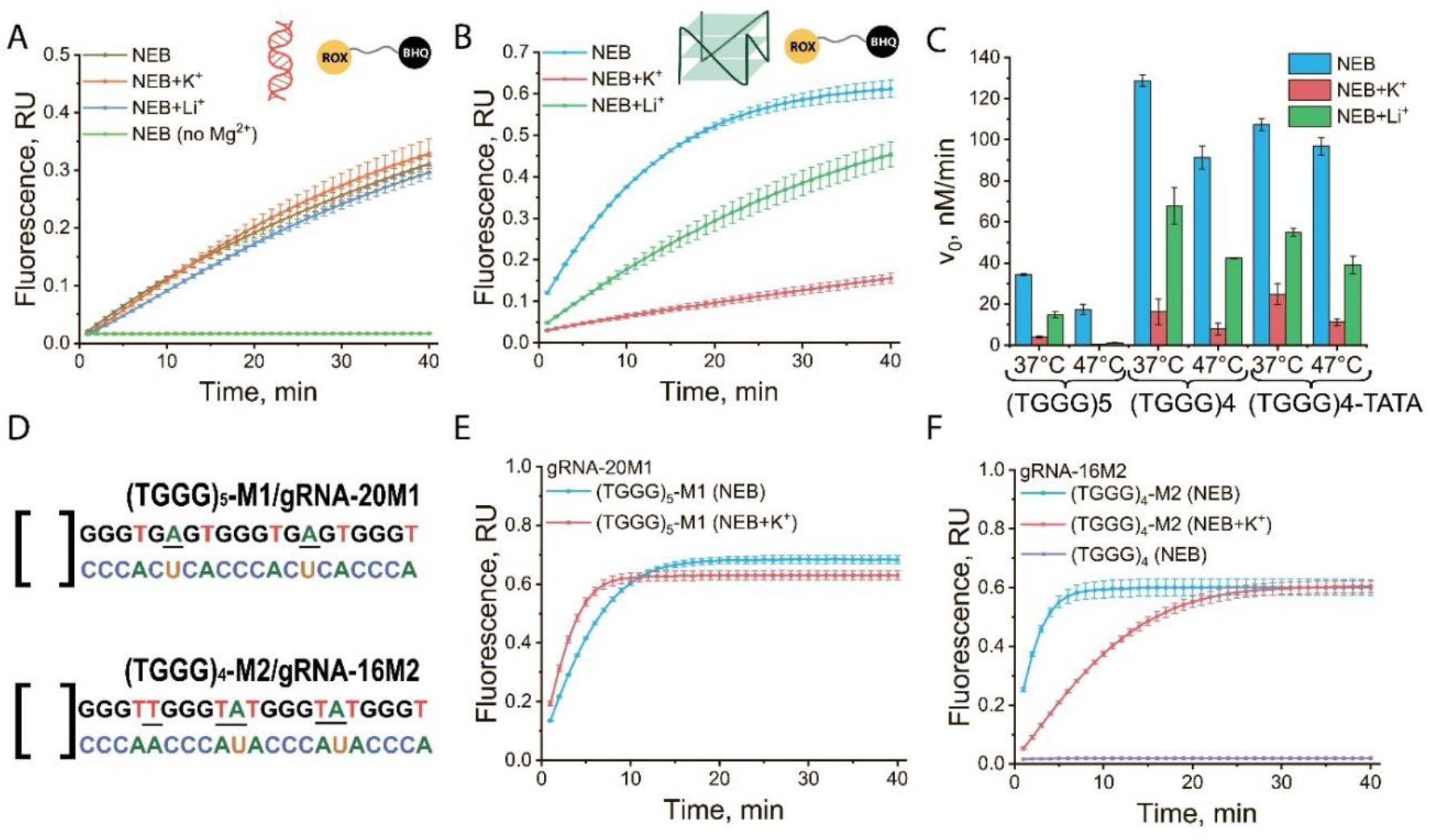
G4 fold stabilization suppresses Cas12a *cis*-activation. (A) Fluorescence curves (LbCas12a, 37 °C) for the dsDNA target (503-bp *C. sepedonicus* amplicon) in NEB, with 50 mM KCl, with 50 mM LiCl or without MgCl_2_. (B) Fluorescence curves for the *cis*-target (TGGG)_5_ in NEB and with 50 mM KCl or 50 mM LiCl (LbCas12a, 37 °C). (C) Initial velocities V_0_ for (TGGG)_5_, (TGGG)_4_ and (TGGG)_4_-TATA in NEB and with 50 mM KCl or 50 mM LiCl at 37 and 47 °C. (D) Sequences of the mutants (TGGG)_5_-M1 (gRNA-20M1; disrupted core) and (TGGG)_4_-M2 (gRNA-16M2; retained quadruplex with elongated loops). (E) Fluorescence curves for (TGGG)_5_-M1 in NEB and with 50 mM KCl (LbCas12a, gRNA-20M1, 37 °C). (F) Fluorescence curves for (TGGG)_4_-M2 in NEB and with 50 mM KCl, and for (TGGG)_4_ in NEB with gRNA-16M2 (LbCas12a, 37 °C). Data are mean ± SD (n = 3).

By contrast, folded G4 *cis*-DNA-targets showed salt-dependent activation. For (TGGG)_5_ at 37 °C, 50 mM KCl strongly reduced the initial rate of LbCas12a activation, whereas 50 mM LiCl produced an intermediate reduction (Fig. 4B). V_0_ values for (TGGG)_5_, (TGGG)_4_ and (TGGG)_4_-TATA at 37 and 47 °C showed that K^+^ produced the strongest suppression across the compact scaffolds (Fig. 4C; full kinetic series for LbCas12a and AsCas12a in SI, Figure S11). The decrease was approximately 8-10-fold for (TGGG)_5_ and (TGGG)_4_ at 37 °C and exceeded one order of magnitude for these scaffolds at 47 °C. For (TGGG)_4_-TATA, K^+^ suppression was smaller but still substantial, approximately 4-5-fold at 37 °C and close to one order of magnitude at 47 °C. Li^+^ generally reduced V_0_ less strongly than K^+^, with the weakest Li^+^ response observed for (TGGG)_5_ at 47 °C. Thus, cation effects on *cis*-activation followed the structural stability series measured by CD and melting: stronger G4 stabilization corresponded to lower activation rates, whereas the *cis*-dsDNA-target control remained active under the same salt conditions.

Mutant analysis further supported this interpretation. The core-disrupting mutant (TGGG)_5_-M1 and the fold-preserving long-loop mutant (TGGG)_4_-M2 (Fig. 4D) were assayed with their cognate gRNAs, gRNA-20M1 and gRNA-16M2. For (TGGG)_5_-M1, K^+^ did not reduce activation after disruption of the G4 core (Fig. 4E). For (TGGG)_4_-M2, which retained G4 folding but contained elongated loops, K^+^ reduced the initial activation rate (Fig. 4F), although less strongly than for the short-loop (TGGG)_4_ scaffold (SI, Figure S11B). A control reaction with (TGGG)_4_ and non-cognate gRNA-16M2 remained at background. Thus, K^+^ suppression requires a folded G4 core, and its magnitude depends on target architecture. These data support a model in which stabilization of the folded *cis*-target limits formation of the gRNA–target hybrid required for Cas12a activation.

These results indicated that Cas12a *cis*-activation was linked to G4 stability: stabilization of the folded target slowed activation, whereas core disruption removed K^+^ sensitivity. This pattern suggested that recognition involved a conformational transition of the G4 *cis*-target from the folded to the unfolded state. The transition was then examined directly using terminal FRET donor and acceptor labels.

### FRET and electrophoretic detection of G4 unfolding and *cis*-target cleavage

The results above suggested that interaction of a compact parallel G4 *cis*-target with the gRNA–Cas12a complex involved G4 unfolding. This conformational transition was monitored directly using (TGGG)_5_ derivatives carrying terminal FRET donor and acceptor labels: AF555-(TGGG)_5_-AF647, FAM-(TGGG)_5_-TAMRA, Coumarin343-(TGGG)_5_-FAM, Cy3-(TGGG)_5_-Cy5 and JOE-(TGGG)_5_-Cy5 (SI, Table S1). In compact folded (TGGG)_5_, the two termini are positioned close enough for donor-sensitized acceptor emission. Hybridization with complementary (CCCA)_5_ or cognate gRNA, and recognition by the gRNA– Cas12a complex, would increase the donor-acceptor distance and reduces the FRET readout (Fig. 5A).

**Figure 5.**
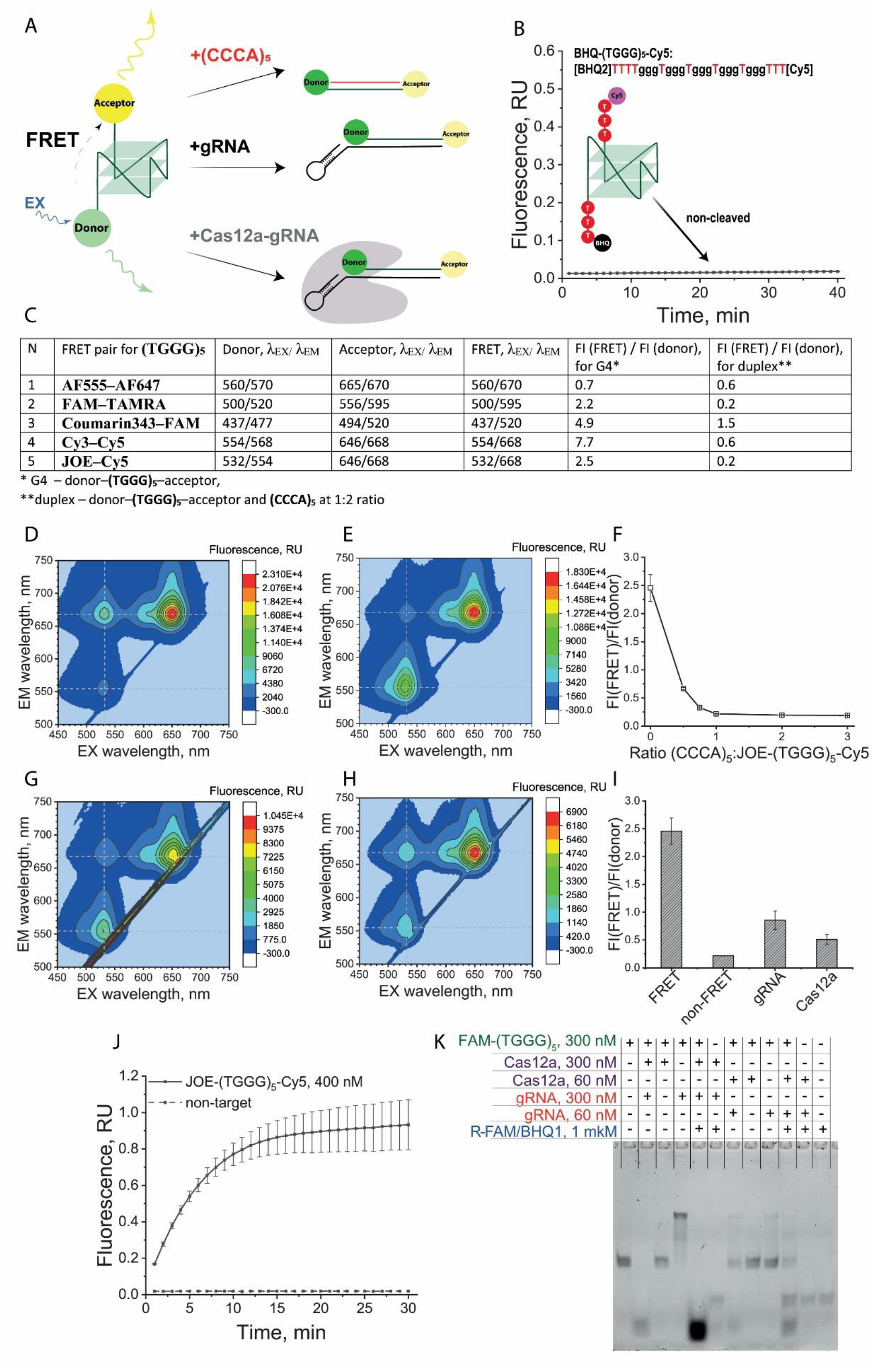
Direct monitoring of G4 unfolding and *cis*-target cleavage. (A) FRET scheme: donor and acceptor labels at the 5′/3′ ends of (TGGG)_5_ are close in the folded quadruplex; duplex formation with (CCCA)_5_, gRNA binding or gRNA–Cas12a recognition separates the labels. (B) Cy5 fluorescence curve (AsCas12a, 37 °C) for the BHQ-(TGGG)_5_-Cy5 construct with TTTT/TTT spacers. (C) Donor/acceptor pairs for (TGGG)_5_: excitation/emission wavelengths and FI(FRET)/FI(donor) ratio for folded G4 and the duplex with (CCCA)_5_ at a 1:2 ratio. (D) 3D fluorescence spectrum of folded JOE-(TGGG)_5_-Cy5. (E) The same construct after duplex formation with (CCCA)_5_. (F) FI(FRET)/FI(donor) (532-nm excitation) versus the (CCCA)_5_:JOE-(TGGG)_5_-Cy5 ratio. (G) 3D spectrum of JOE-(TGGG)_5_-Cy5 after incubation with gRNA-AsCas12a. (H) The same with gRNA alone. (I) FI(FRET)/FI(donor) (532-nm excitation) for folded G4, duplex, gRNA-treated construct and Cas12a/gRNA-treated construct. (J) Fluorescence curves (AsCas12a, 37 °C) showing R-FAM/BHQ1 reporter cleavage after incubation of JOE-(TGGG)_5_-Cy5 (400 nM) with gRNA–Cas12a. (K) Denaturing PAGE (7 M urea, in-gel FAM detection) of FAM-(TGGG)_5_ *cis*-target (400 nM) in reactions containing combinations of Cas12a (300 or 60 nM), gRNA (300 or 60 nM) and R-FAM/BHQ1 reporter (1 µM). Data are mean ± SD (n = 3).

Short TTTT/TTT spacers were inserted between the G4 core and terminal labels to reduce local dye– guanine interactions, including guanine-mediated fluorescence quenching, and to minimize direct perturbation of G4 folding by the dyes. The spacer-containing architecture was first evaluated with the single-fluorophore/dark-quencher construct BHQ-(TGGG)_5_-Cy5, initially in the role of a labelled *trans*-target reporter. In an AsCas12a collateral-cleavage assay activated by the 503-bp *cis*-dsDNA-target, this construct did not show a Cy5 fluorescence increase (Fig. 5B), indicating that the labelled spacer-containing construct retained a folded G4 state resistant to *trans*-cleavage. By contrast, addition of complementary (CCCA)_5_ separated the quencher and fluorophore and increased Cy5 fluorescence (SI, Figure S12A,B,E). Thus, the labels responded to strand separation, but the gRNA–Cas12a-induced change was small and depended on labelled *cis*-DNA-target concentration (SI, Figure S12C,D). Subsequent conformational analysis therefore used FRET donor/acceptor constructs and the ratio of donor-sensitized acceptor emission to donor emission, FI(FRET)/FI(donor).

Excitation-emission maps first defined the excitation/emission maxima for donor fluorescence, acceptor fluorescence and FRET, corresponding to donor excitation with acceptor emission (Fig. 5C-E; SI, Figures S19–S22). Fluorescence intensities (FI) were then measured at the selected wavelengths. FI(FRET)/FI(donor), calculated as acceptor-emission intensity under donor excitation divided by donor-emission intensity under donor excitation, was used to compare folded and duplex states. The complement ratio required to generate a low-FRET duplex reference was determined using JOE-(TGGG)_5_-Cy5. Folded JOE-(TGGG)_5_-Cy5 showed a clear Cy5 emission signal upon JOE excitation, with maxima at 532-nm excitation and 668-nm emission (Fig. 5D). Addition of complementary (CCCA)_5_ reduced this signal and increased JOE emission at 554 nm under 532-nm excitation (Fig. 5E). Titration of (CCCA)_5_ from 0.5 to 3 equivalents relative to JOE-(TGGG)_5_-Cy5 progressively decreased FI(FRET)/FI (donor), with minimal values reached at 2:1 and 3:1 complement:G4 ratios (Fig. 5F; 3D spectra at different dilutions and molar ratios in SI, Figures S13–S18). Thus, a twofold excess of complement was sufficient to define the low-FRET duplex reference.

Using the 2:1 complement:G4 duplex reference, the donor/acceptor-labelled constructs were compared for folded-to-duplex separation. Three pairs showed the largest changes in FI(FRET)/FI(donor): Cy3–Cy5, JOE–Cy5 and FAM–TAMRA, with 12.8-, 12.5- and 11-fold differences, respectively. AF555–AF647 and Coumarin343–FAM gave smaller differences and were not analyzed further (Fig. 5C; SI, Figures S19– S22). The three selected constructs were then tested after incubation with gRNA–AsCas12a. All showed lower FI(FRET)/FI(donor), consistent with G4 unfolding during recognition by the gRNA–AsCas12a complex (Fig. 5G-I; SI, Figures S20, S21 and S24). The largest decrease was observed for JOE-(TGGG)_5_-Cy5. For this construct, gRNA-20 alone also reduced FI(FRET)/FI(donor), indicating partial unfolding during gRNA–target hybrid formation; addition of AsCas12a lowered the ratio further toward the low-FRET duplex reference (Fig. 5G-I; SI, Figures S23 and S24). After incubation of JOE-(TGGG)_5_-Cy5 with gRNA-20/AsCas12a, addition of R-FAM/BHQ1 produced an increasing reporter signal (Fig. 5J), confirming that recognition of the labelled G4 *cis*-target activated collateral *trans*-cleavage.

Denaturing PAGE was used to determine whether recognition of the G4 as a *cis*-DNA-target was accompanied only by unfolding or also by cleavage of the target strand. Reactions were assembled with combinations of the *cis*-DNA-target bearing a single FAM label on an extended 5′ end, FAM-(TGGG)_5_, Cas12a, gRNA and the R-FAM/BHQ1 reporter. Products were separated under denaturing conditions in 7 M urea (Fig. 5K). In samples where the CRISPR–Cas12a reaction could not proceed because a key component was missing, a single FAM-positive band corresponding to full-length, non-cleaved FAM-(TGGG)_5_ was observed. In samples containing the required components, an additional faster-migrating band corresponding to a shorter FAM-labelled fragment appeared. Because urea abolishes secondary structure, this mobility shift reflects scission of the phosphodiester backbone rather than a conformational change. Electrophoresis therefore provided evidence that folded (TGGG)_5_ is cleaved during recognition by the gRNA–Cas12a complex. The effect depended on gRNA/Cas12a concentration: the 300 nM gRNA / 300 nM Cas12a complex gave a more intense cleaved FAM-(TGGG)_5_ band than the 60 nM gRNA / 60 nM Cas12a complex.

Therefore, FRET and electrophoretic analyses explain how a compact parallel G4 acts as a *cis*-target for the gRNA–Cas12a complex: recognition proceeds through unfolding of the quadruplex, after which the strand is cleaved as a *cis*-target, whereas the same folded motif resists collateral *trans*-cleavage. The following experiments examined whether this folding-encoded asymmetry was specific to compact parallel G4s or also occurred in structurally distinct antiparallel quadruplexes.

### Antiparallel G4s show folding-dependent suppression without full *trans*-resistance

The folding-defined asymmetry observed for compact parallel (TGGG)n was next evaluated with structurally distinct G4 scaffolds. The set included two sequences assigned as antiparallel in previous studies: the quadruplex aptamer RE21 [30] and the anti-ochratoxin A aptamer aptOTA [31], together with the truncated aptOTA variant aptOTA-cut (Fig. 6A). Under the present buffer conditions, CD confirmed an antiparallel G4 signature for aptOTA-cut, with a positive band near 295 nm and a negative band near 265 nm at 25 and 37 °C (Fig. 6F), and a similar spectrum was observed for aptOTA (SI, Figure S25B). RE21 did not show a defined G4 spectrum in NEBuffer 2.1 (SI, Figure S25A).

**Figure 6.**
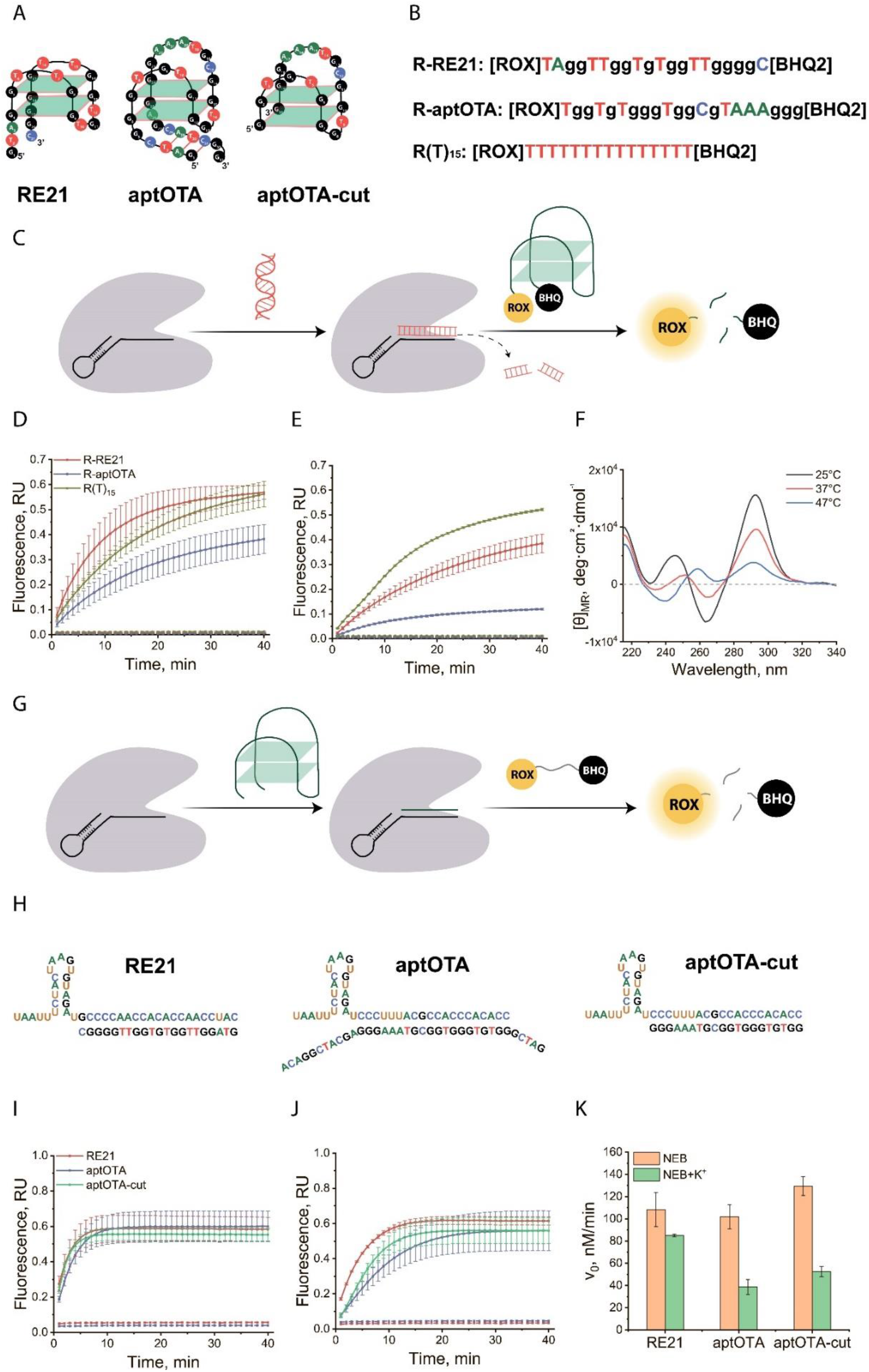
Antiparallel G4s show partial, stability-dependent effects on Cas12a. (A) Schematic representations of RE21, aptOTA and aptOTA-cut. (B) Sequences of the reporters R-RE21, R-aptOTA and standard R(T)_15_. (C) Scheme for testing an antiparallel G4 reporter as a collateral-cleavage substrate. (D) Fluorescence curves for R-RE21 and R-aptOTA compared with R(T)_15_ (LbCas12a, 37 °C) in NEB and (E) with 50 mM KCl; dashed lines, no-target controls with water. (F) CD spectra of aptOTA-cut at 25, 37 and 47 °C in NEB. (G) Scheme for testing an antiparallel G4 as a *cis*-target. (H) Cognate gRNA/ssDNA target pairs for RE21, aptOTA and aptOTA-cut. (I) Fluorescence curves for RE21, aptOTA and aptOTA-cut *cis*-targets (LbCas12a, 37 °C, R(T)_15_ readout) in NEB and (J) with 50 mM KCl; dashed lines, water controls. (K) Initial velocities V_0_ for RE21, aptOTA and aptOTA-cut in NEB and with 50 mM KCl at 37 °C. Data are mean ± SD (n = 3).

Antiparallel G4s were first tested as labelled *trans*-target reporters using ROX/BHQ-labelled R-RE21 and R-aptOTA (Fig. 6B,C). Both reporters were cleaved by activated LbCas12a in NEBuffer 2.1 (Fig. 6D). The R-RE21 signal was close to that of the linear reference R(T)_15_, whereas R-aptOTA produced a lower but clearly detectable signal. This behaviour contrasts with compact parallel (TGGG)n reporters, which were resistant to collateral *trans*-cleavage. Addition of 50 mM KCl reduced cleavage of R-RE21 and R-aptOTA (Fig. 6E), while the same K^+^ concentration did not reduce cleavage of the linear R(T)_15_ reporter. A similar K^+^-dependent reduction was observed with AsCas12a (SI, Figure S26). Thus, antiparallel G4 reporters remained accessible to collateral cleavage, but K^+^-dependent stabilization partially protected them.

The same sequences were then tested as guide-complementary *cis*-targets (scheme, Fig. 6G; gRNA/target pairs, Fig. 6H). RE21, aptOTA and aptOTA-cut activated LbCas12a in NEBuffer 2.1 (Fig. 6I). Addition of 50 mM KCl reduced activation to different extents (Fig. 6J). V_0_ comparison at 37 °C showed little K^+^ effect for RE21, whereas aptOTA and aptOTA-cut showed approximately two-fold or stronger reduction (Fig. 6K). This pattern agrees with the CD data: RE21 lacked a defined G4 fold in NEBuffer 2.1, whereas aptOTA and aptOTA-cut formed antiparallel G4s whose stabilization by K^+^ reduced *cis*-activation. Similar trends were observed at 47 °C (SI, Figure S27). Thus, as with the parallel motifs, stabilization of a folded G4 *cis*-DNA-target impaired Cas12a activation, consistent with less efficient opening of the folded target.

These data extend the folding-dependent control observed for compact parallel G4s to antiparallel scaffolds, but with a weaker effect on collateral cleavage. Antiparallel G4s retained *cis*-activation and showed K^+^-dependent suppression of both *cis*-activation and *trans*-cleavage, whereas compact parallel (TGGG)n combined retained *cis*-activation with complete *trans*-resistance.

## Discussion

This study identifies G4 folding as a structural determinant that separates the two Cas12a activities used in nucleic-acid diagnostics. Compact parallel (TGGG)n G4s activated LbCas12a and AsCas12a as guide-complementary *cis*-targets, but the same folded motifs did not generate collateral-cleavage signals as labelled *trans*-target reporters. Core disruption and G-rich non-G4 controls restored *trans*-cleavage, demonstrating that resistance depends on quadruplex architecture. K^+^ stabilization reduced *cis*-activation rates by approximately 8-10-fold at 37 °C and by more than one order of magnitude at 47 °C for the most stable parallel G4 constructs. FRET and denaturing PAGE showed that productive *cis*-recognition involves G4 unfolding followed by target-strand cleavage. Thus, the same compact ssDNA fold can support Cas12a activation while preventing efficient collateral *trans*-cleavage.

The differential response is consistent with distinct requirements for *cis*-recognition and *trans*-cleavage. In the *cis* position, formation of the guide-target hybrid requires opening of the folded ssDNA target; once formed, the hybrid can stabilize the recognized state and permit nuclease activation [8,9]. In the *trans* position, activated Cas12a has no complementary guide interaction with the reporter, and cleavage therefore depends on local accessibility of the ssDNA backbone to the nonspecific active site. Compact parallel G4s with short loops limit this accessibility, as indicated by their resistance to collateral *trans*-cleavage and Exonuclease I, and by restored cleavage after disruption of the G4 core. This explains how Cas12a can remodel a folded *cis*-target yet cleave the same architecture inefficiently as a *trans*-target.

These results extend earlier Cas12a studies that used G4 motifs mainly as structure-responsive reporters or reporter-derived *trans*-targets [4,5,16–19]. Those studies established that G4 folding can slow, tune or suppress *trans*-cleavage under selected conditions. Here, compact parallel (TGGG)n G4s showed complete *trans-*resistance under the tested conditions. This distinguishes them from the less compact G4 reporters examined previously and is consistent with the partial, rather than complete, *trans*-protection observed for antiparallel G4s in this study. At the same time, the same folded (TGGG)n scaffold acted as a guide-complementary *cis*-target. Cas9 and dCas9 studies provided the closest precedents by showing that CRISPR guide complexes can interact with G4-containing DNA [20–24]. The present data extend this principle to Cas12a and to single-stranded G4 *cis*-targets, where recognition is linked to G4 unfolding, target-strand cleavage and activation of collateral *trans*-cleavage. Thus, compact G4 folding creates a position-dependent response in which the motif is unfolded and cleaved during *cis*-recognition but resists nonspecific collateral *trans*-cleavage as a reporter.

The comparison across sequence variants and topologies indicates that architecture and fold stability, rather than guanine content alone, determine the Cas12a response. G-rich non-G4 controls were cleaved, and disruption of the (TGGG)n core eliminated *trans*-resistance and K^+^-dependent suppression of *cis*-activation. Antiparallel RE21 and aptOTA-family G4s behaved differently: they retained *cis*-activation but showed only partial K^+^-dependent protection in *trans*. Compact parallel (TGGG)n therefore represents the strongest limit of a graded folding-dependent effect, whereas less compact or less stable G4 topologies produce partial suppression.

These findings have practical implications for CRISPR–Cas12a diagnostics. Secondary structure in *cis*-targets and *trans*-target reporters is a determinant of both activation kinetics and reporter cleavage. Unintended G4 folding in G-rich targets or reporters may therefore change assay readouts even when the primary sequence appears suitable. Conversely, compact G4 modules can be used deliberately as condition-tunable reporter elements. Because G4 stability can be changed by ions, sequence and temperature, structural gating provides a direct way to tune diagnostic signal generation without changing the nuclease or guide RNA.

## Funding

The study was supported by the Russian Science Foundation (No. 25-16-00246).

## Acknowledgements

During the preparation of this manuscript, the authors used an AI-based tool to assist with language editing. The authors reviewed and edited the text as needed and take full responsibility for the content of the publication.

## Data Availability

All data supporting the findings of this study are available within the article and its Supplementary Information.

## Notes

### Competing Interest Statement

The authors have declared no competing interest.

