## Supplementary material for "G-quadruplex folding uncouples cis-activation from collateral trans-cleavage by Cas12a": SI

**Table S1.** Sequences of oligonucleotide used in this research

| No | Name | Sequence 5’-3’ | Purpose |
|---|---|---|---|
| 1 | R$(TGGG)_4$ | [ROX]TgggTgggTgggTggg[BHQ2] | Trans-targets, fluorophore-quencher reporters |
| 2 | R$(TGGG)_5$ | [ROX]TgggTgggTgggTgggTggg[BHQ2] | |
| 3 | R$(TGGG)_4$-TATA | [ROX]TgggTgggTgggTgggTATA[BHQ2] | |
| 4 | R1 | [ROX]TggAAgggAAgggTTT[BHQ2] | |
| 5 | R2 | [ROX]TggAAgggAAggg[BHQ2] | |
| 6 | R3 | [ROX]TggAAgggAAgggTTTTT[BHQ2] | |
| 7 | R4 | [ROX]TgTggTggTggTggTgg[BHQ2] | |
| 8 | R$(T)_{15}$ | [ROX]TTTTTTTTTTTTTTT[BHQ2] | |
| 9 | R-FAM/BHQ1 | [FAM]CCCAACGAGAAGCG[BHQ1] | |
| 10 | R$(TGGG)_5$-M1 | [ROX]TgggTg**A**gTgggTg**A**gTggg[BHQ2] | |
| 11 | R-RE21 | [ROX]TAGGTTGGTGTGGTTGGGGC[BHQ] | |
| 12 | R-aptOTA | [ROX]TGGTGTGGGTGGCGTAAAGGG[BHQ] | |
| 13 | $(TGGG)_4$ | TgggTgggTgggTggg | Single-stranded cis-targets |
| 14 | $(TGGG)_5$ | TgggTgggTgggTgggTggg | |
| 15 | $(TGGG)_4$-TATA | TgggTgggTgggTgggTATA | |
| 16 | $(TGGG)_5$-M1 | TgggTg**A**gTgggTg**A**gTggg | |
| 17 | $(TGGG)_4$-M2 | TgggT**AT**gggT**AT**gggT**T**ggg | |
| 18 | RE21 | GTAGGTTGGTGTGGTTGGGGC | |
| 19 | aptOTA | GATCGGGTGTGGGTGGCGTAAAGGGAGCATCGGACA | |
| 20 | aptOTA-cut | GGTGTGGGTGGCGTAAAGGG | |
| 21 | $(TGGG)_2$ | TgggTggg | |
| 22 | $(TGGG)_3$ | TgggTgggTggg | |
| 23 | FAM-$(TGGG)_5$ | [FAM]TTTTTTTgggTgggTgggTgggTggg | Cis-targets for electrophoresis |
| 24 | $(CCCA)_4$ | CCCACCCACCCACCCA | |

| | | | |
|---|---|---|---|
| 25 | $(CCCA)_5$ | CCCACCCACCCACCCACCCA | Sequences complementary to quadruplexes |
| 26 | TATA-$(CCCA)_4$ | TATACCCACCCACCCACCCA | |
| 27 | AF555-$(TGGG)_5$-AF647 | [AF555]TTTTgggTgggTgggTgggTgggTTT[AF647] | FRET constructs |
| 28 | FAM-$(TGGG)_5$-TAMRA | [FAM]TTTTgggTgggTgggTgggTgggTTT[TAMRA] | |
| 29 | Coumarin343-$(TGGG)_5$-FAM | [Coumarin343]TTTTgggTgggTgggTgggTgggTTT[FAM] | |
| 30 | Cy3-$(TGGG)_5$-Cy5 | [Cy3]TTTTgggTgggTgggTgggTgggTTT[Cy5] | |
| 31 | JOE-$(TGGG)_5$-Cy5 | [JOE]TTTTgggTgggTgggTgggTgggTTT[Cy5] | |
| 32 | BHQ-$(TGGG)_5$-Cy5 | [BHQ2]TTTTgggTgggTgggTgggTgggTTT[Cy5] | |
| 33 | gRNA-16 | uaauuucuacuaaguguagauCCCACCCACCCACCCA | gRNA for $(TGGG)_4$ |
| 34 | gRNA-16/4 | uaauuucuacuaaguguagauUAUACCCACCCACCCACCCA | gRNA for $(TGGG)_4$-TATA |
| 35 | gRNA-20 | uaauuucuacuaaguguagauCCCACCCACCCACCCACCCA | gRNA for $(TGGG)_5$ |
| 36 | gRNA-20M1 | uaauuucuacuaaguguagauCCCACUCACCCACUCACCCA | gRNA for $(TGGG)_5$-M1 |
| 37 | gRNA-16M2 | uaauuucuacuaaguguagauCCCAACCCAUACCCAUACCCA | gRNA for $(TGGG)_4$-M2 |
| 38 | gRNA-RE21 | uaauuucuacuaaguguagauGCCCCAACCACACCAACCUAC | gRNA for RE21 |
| 39 | gRNA-aptOTA | uaauuucuacuaaguguagauCCCUUUACGCCACCCACACC | gRNA for aptOTA/ aptOTA-cut |
| 40 | gRNA-Cs-82 | uaauuucuacuaaguguagauGCCUCCCCUAGAGGAGACAGAA | gRNA for DNA-target of *Clavibacter sepedonicus* at 47 °C |
| 41 | gRNA-Cs-48 | uaauuucuacuaaguguagauAAGAUUCUAAGAGCAAACGGUG | gRNA for DNA-target of *Clavibacter sepedonicus* at 37 °C |
| 42 | PCRCs-F | CTCCTTGTGGGGTGGGAAAA | PCR primers to obtain 503-bp fragment of *C. sepedonicus* |
| 43 | PCRCs-R | TACTGAGATGTTTCACTTCCCC | |

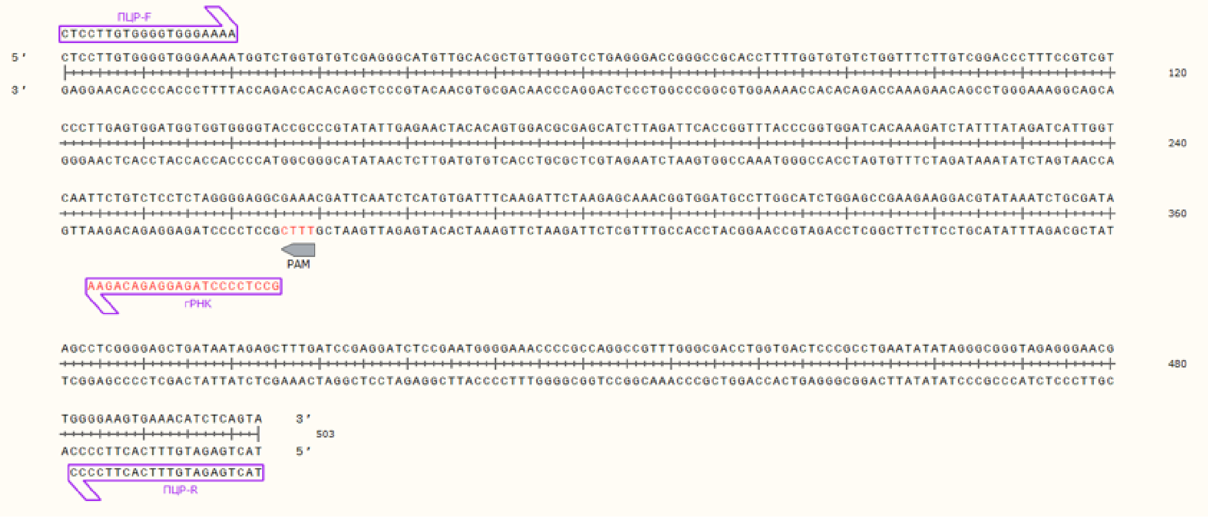


**Figure S1**. Fragment of the 16S–23S locus from the reference genome showing the location of the PAM, the PCR primers and the gRNA Cs-82 used for the CRISPR/Cas12a reaction (image from SnapGene Viewer, GSL Biotech LLC, USA).

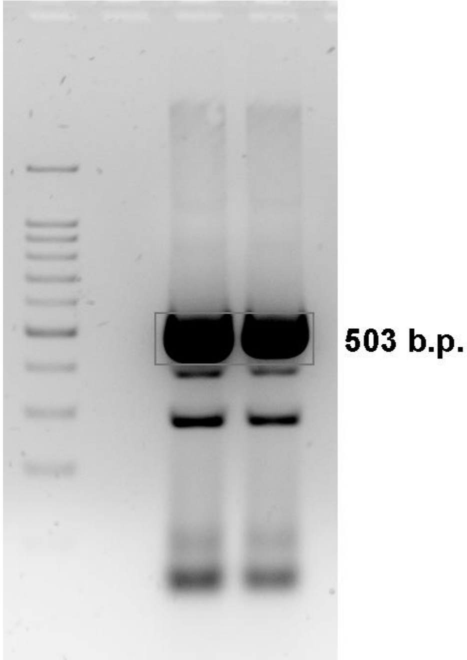


**Figure S2**. UV fluorescence of the PCR product obtained from *C. sepedonicus* total DNA after electrophoresis in a 2% agarose gel stained with ethidium bromide; left, 100 bp+ DNA marker; right, PCR product in two replicates.

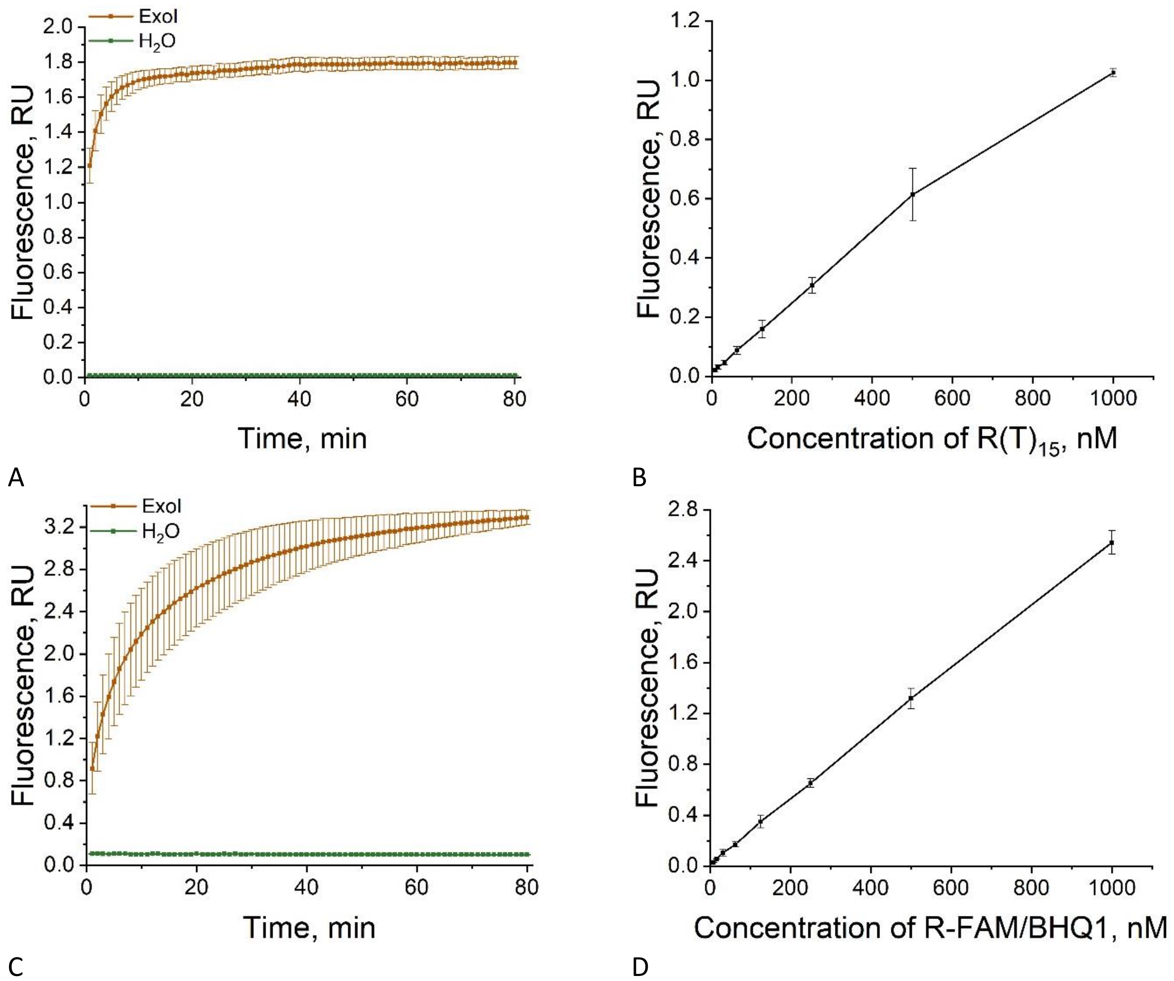


**Figure S3**. Fluorescence curves from cleavage of 2 μM reporter $R(T)_{15}$ (A) and R-FAM/BHQ1 (C) by Exonuclease I (negative control, water). Concentration dependence of the fluorescence signal for cleaved $R(T)_{15}$ (B) and R-FAM/BHQ1 (D) after stopping the reaction with EDTA. Linear fit for $R(T)_{15}$: Fl = 0.00101·c + 0.01599; for R-FAM/BHQ1: Fl = 0.00258·c + 0.01331.

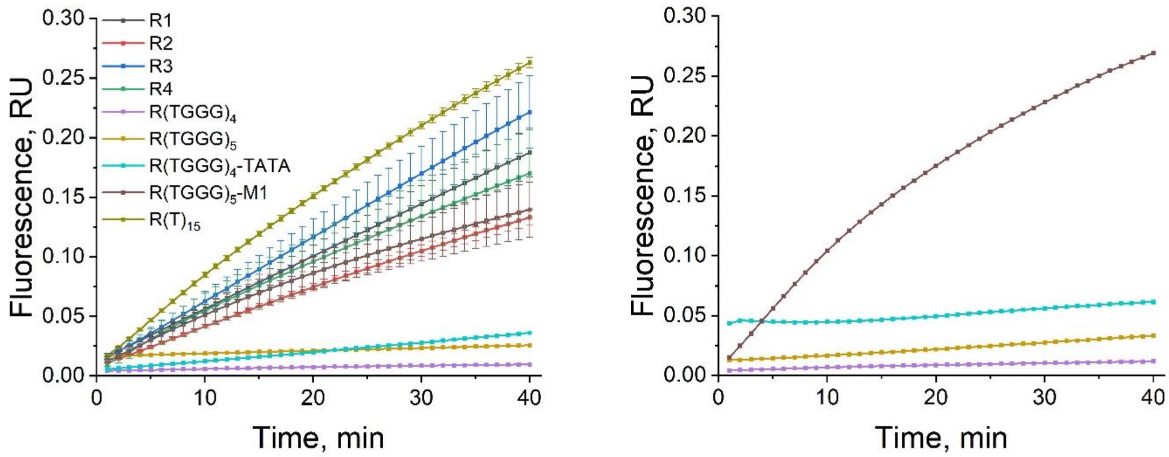


**Figure S4**. Fluorescence curves showing the role of the reporters R1, R2, R3, R4, R(TGGG)$_4$, R(TGGG)$_5$, R(TGGG)$_4$-TATA and R(TGGG)$_5$-M1 as *trans*-targets in CRISPR/LbCas12a at 47 °C, compared with the standard reporter R(T)$_{15}$ (A). Fluorescence curves for R(TGGG)$_4$, R(TGGG)$_5$, R(TGGG)$_4$-TATA and R(TGGG)$_5$-M1 as *trans*-targets in CRISPR/AsCas12a at 47 °C (B).

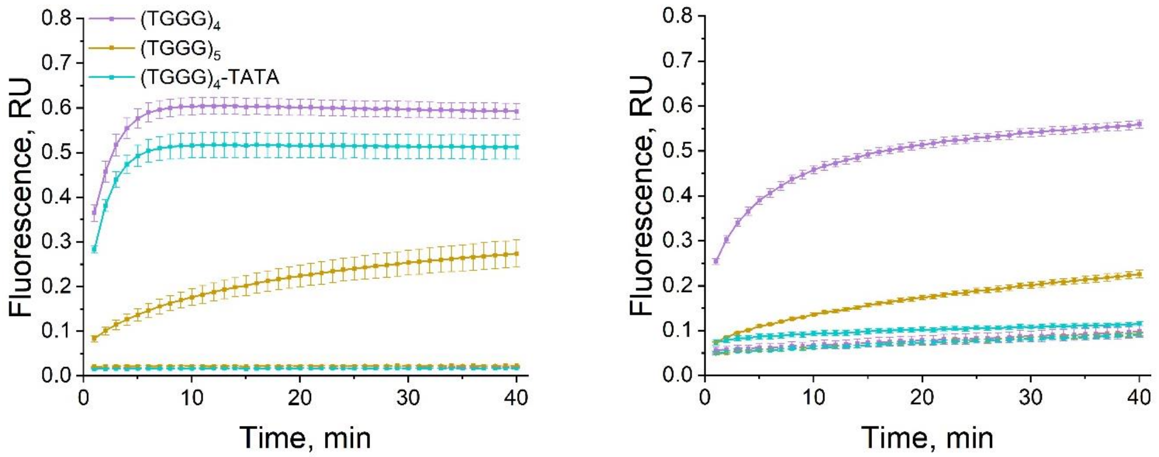


**Figure S5**. Fluorescence curves showing the role of the G-quadruplexes (TGGG)$_4$, (TGGG)$_5$ and (TGGG)$_4$-TATA as *cis*-targets in CRISPR/LbCas12a at 47 °C with R(T)$_{15}$ readout; dashed lines, no-target controls (water) (A). The same in CRISPR/AsCas12a at 47 °C with R-FAM/BHQ1 readout; dashed lines, no-target controls (water) (B).

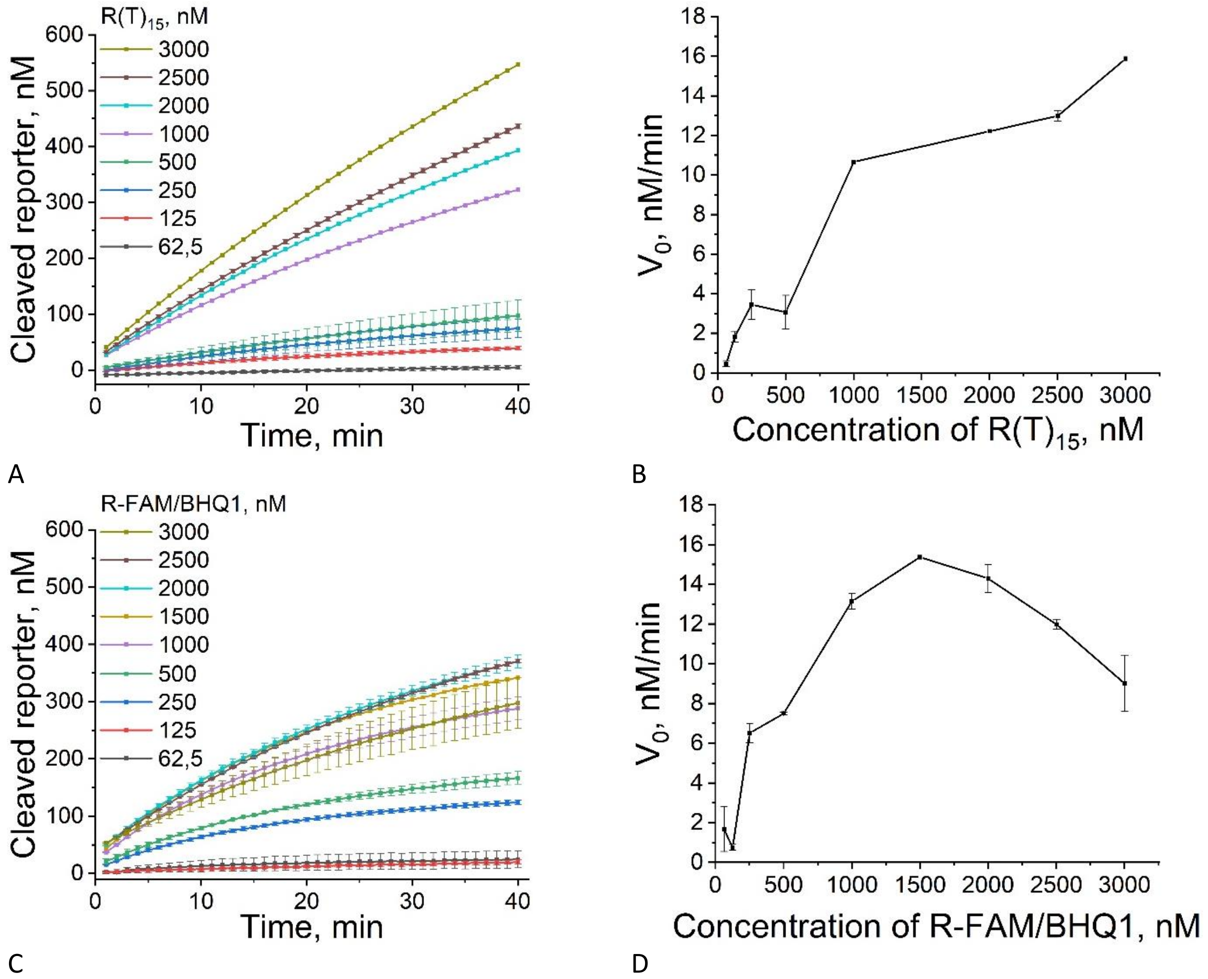


**Figure S6**. Fluorescence curves from the CRISPR/LbCas12a reaction at 37 °C for 42 nM $(TGGG)_5$ (gRNA:LbCas12a = 1:1 = 5.5 nM) with R(T)$_{15}$ read-out at various reporter concentrations (A). Reaction rate versus R(T)$_{15}$ concentration (over the first 4 min), linear coordinates; Michaelis–Menten constant $K_M = 8.2\times10^{-7}$ M, $k_{cat} = 2.9$ min$^{-1}$ (0.05 s$^{-1}$), $k_{cat}/K_M = 5.87\times10^4$ M$^{-1}$ s$^{-1}$ (B). The same for CRISPR/AsCas12a with R-FAM/BHQ1 read-out (C) and the corresponding rate plot; $K_M = 5.22\times10^{-7}$ M, $k_{cat} = 2.79$ min$^{-1}$ (0.05 s$^{-1}$), $k_{cat}/K_M = 8.92\times10^4$ M$^{-1}$ s$^{-1}$ (D).

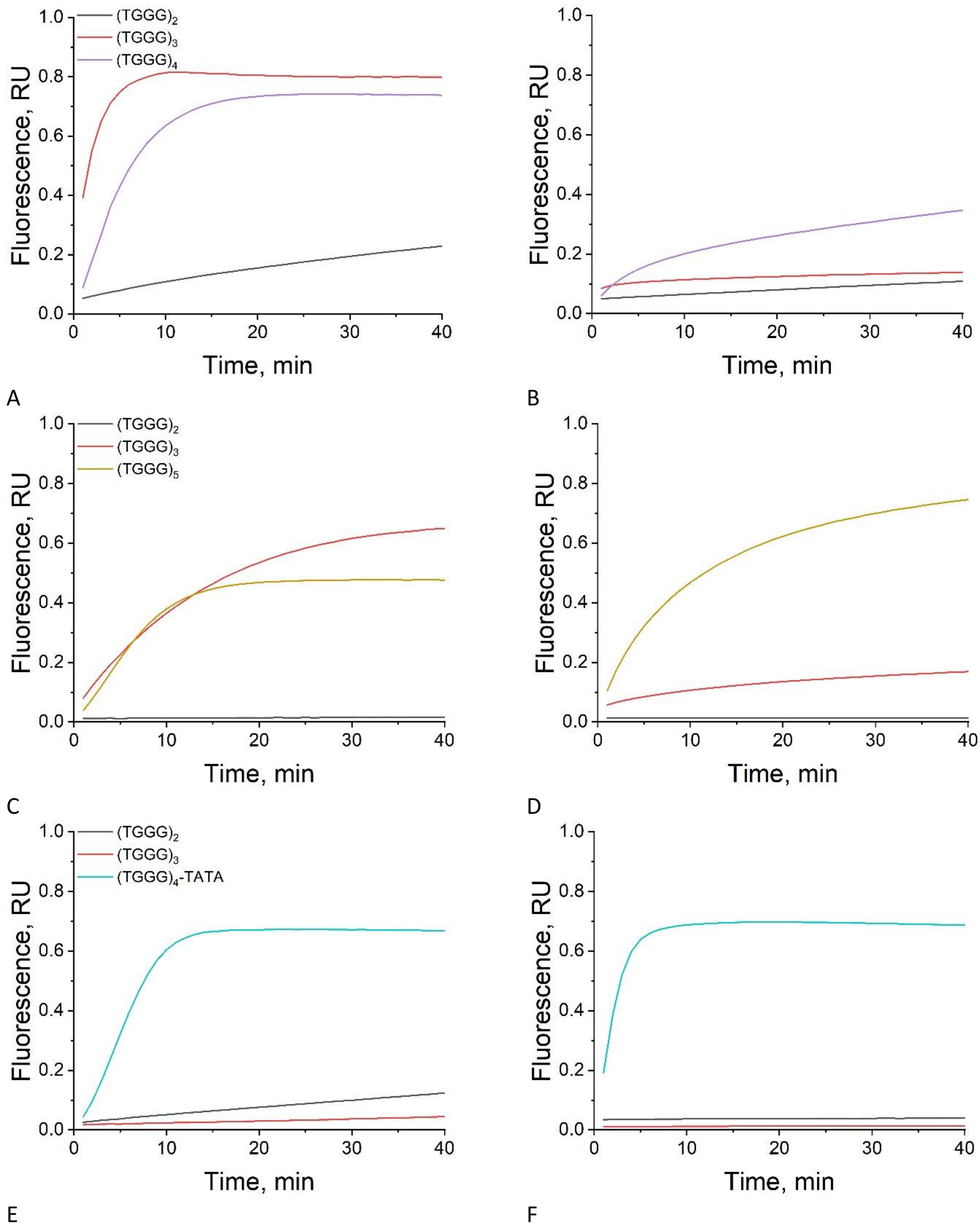


**Figure S7**. Fluorescence curves from the CRISPR/LbCas12a reaction (R(T)$_{15}$ readout) for the shortened sequences (TGGG)$_2$ and (TGGG)$_3$ and the main targets: (TGGG)$_4$ with gRNA-16 at 37 °C (A) and 47 °C (B), (TGGG)$_5$ with gRNA-20 at 37 °C (C) and 47 °C (D), and (TGGG)$_4$-TATA with gRNA-16/4 at 37 °C (E) and 47 °C (F).

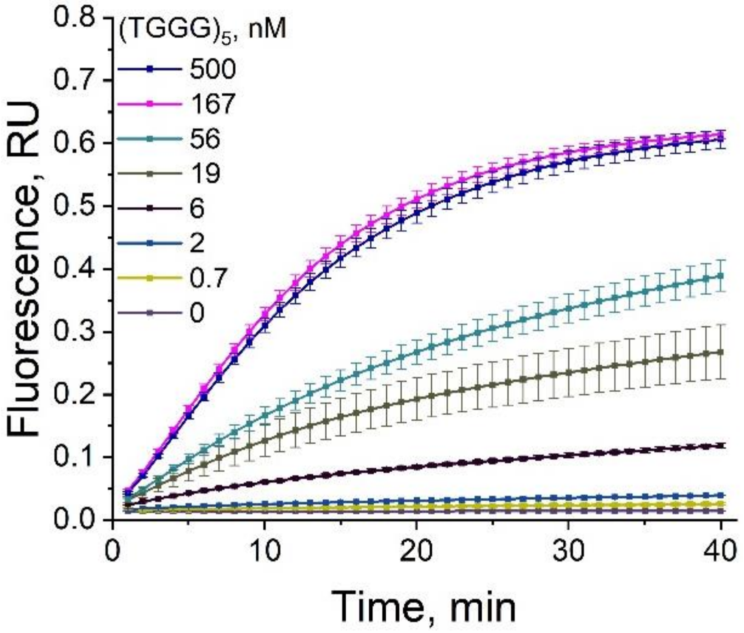


**Figure S8**. Fluorescence curves from the CRISPR/LbCas12a reaction ($R(T)_{15}$ readout) for several concentrations of $(TGGG)_5$ with gRNA-20 at 37 °C.

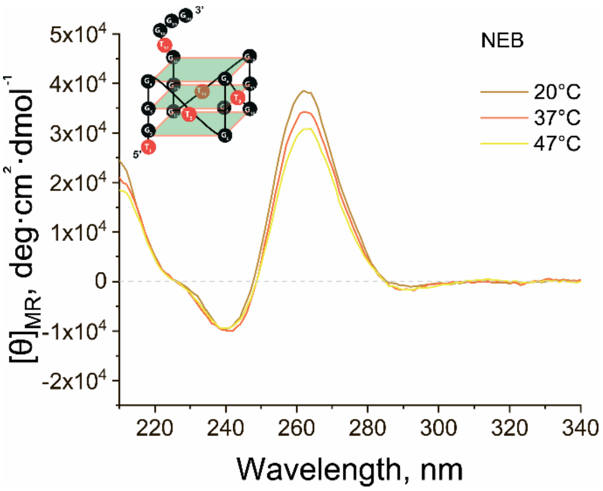
NEB
20°C
37°C
47°C
[θ]MR, deg·cm2·dmol-1
Wavelength, nm

A

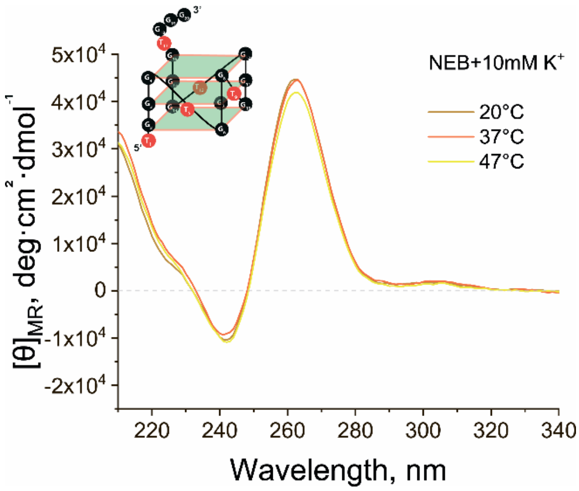
NEB+10mM K+
20°C
37°C
47°C
[θ]MR, deg·cm2·dmol-1
Wavelength, nm

B

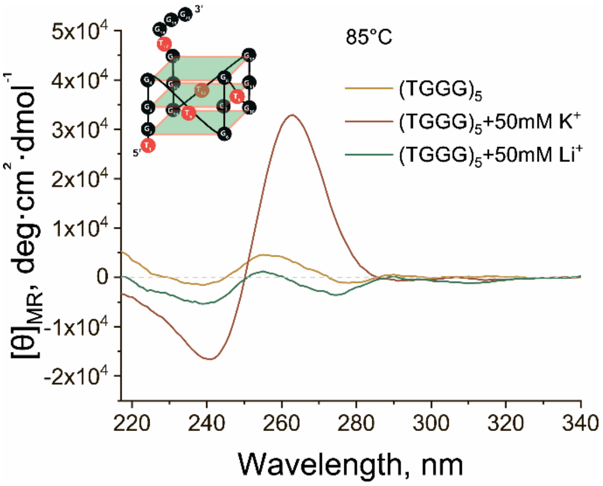
85°C
(TGGG)5
(TGGG)5+50mM K+
(TGGG)5+50mM Li+
[θ]MR, deg·cm2·dmol-1
Wavelength, nm

C

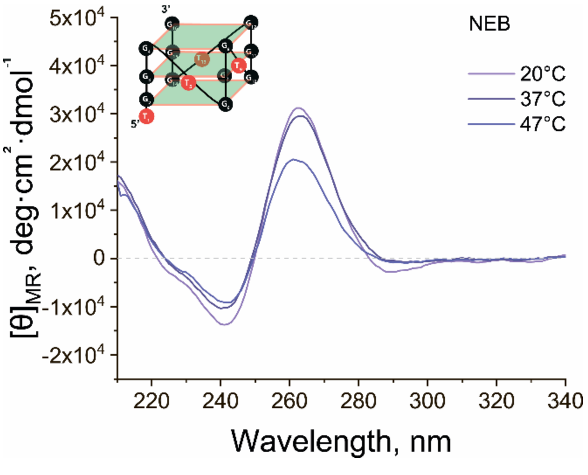
NEB
20°C
37°C
47°C
[θ]MR, deg·cm2·dmol-1
Wavelength, nm

D

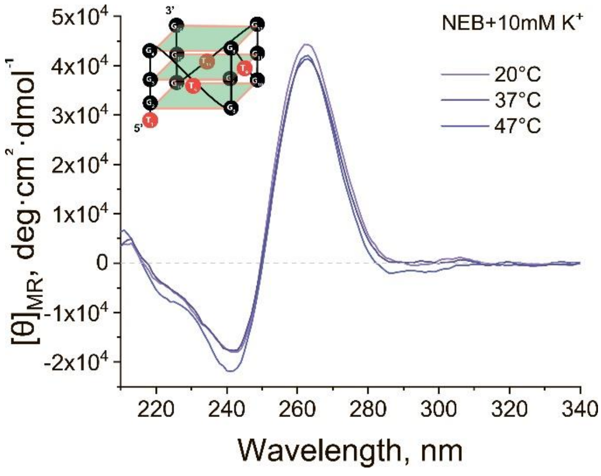
NEB+10mM K+
20°C
37°C
47°C
[θ]MR, deg·cm2·dmol-1
Wavelength, nm

E

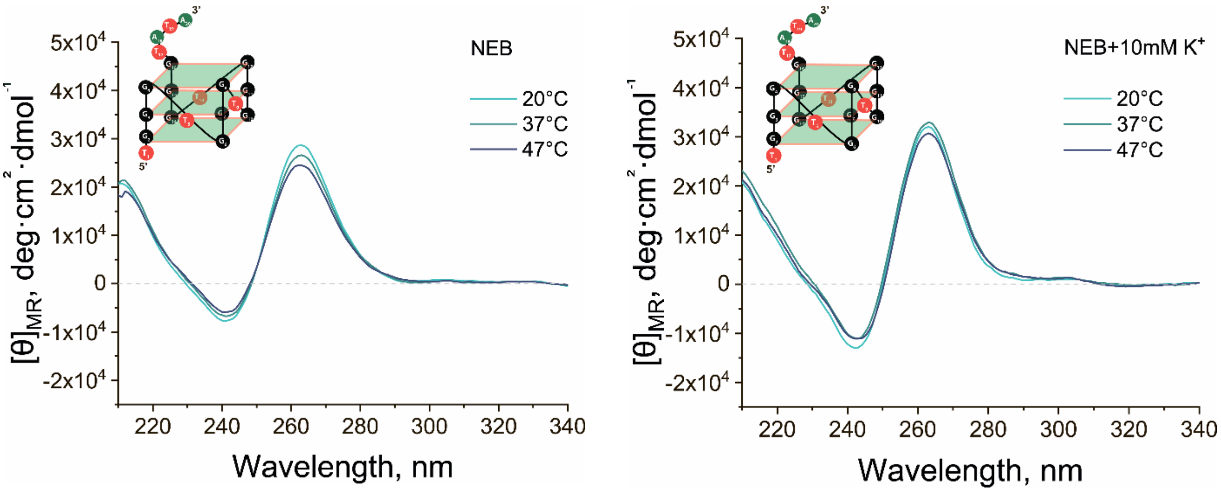


F G

**Figure S9**. CD spectra at 20, 37 and 47 °C for $(TGGG)_5$ in NEB buffer (A) and with 10 mM KCl (B); CD at 85 °C for $(TGGG)_5$ in NEB, with 50 mM KCl and with 50 mM LiCl (C); CD spectra at 20, 37 and 47 °C for $(TGGG)_4$ in NEB (D) and with 10 mM KCl (E), and for $(TGGG)_4$-TATA in NEB (F) and with 10 mM KCl (G).

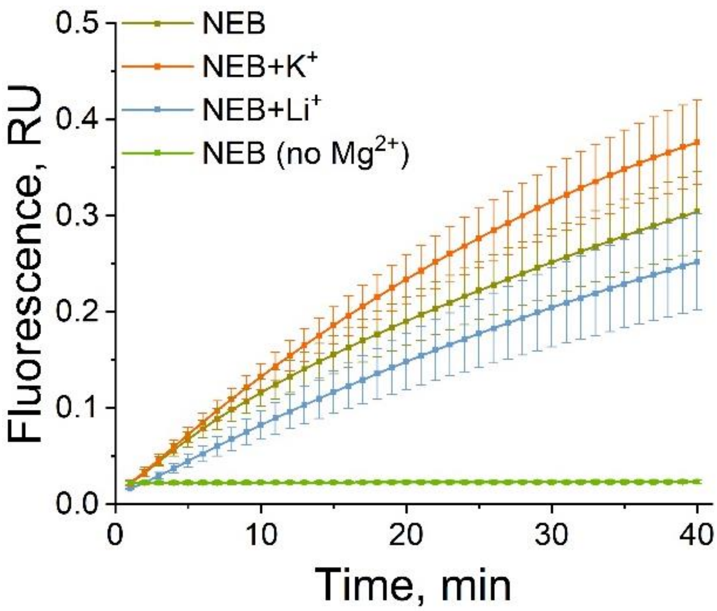


**Figure S10**. Fluorescence curves from the CRISPR/LbCas12a reaction at 47 °C for the dsDNA target (*C. sepedonicus* PCR amplicon; solid lines) in NEB and with 50 mM KCl, 50 mM LiCl or without $MgCl_2$; dashed line, no-target control (water).

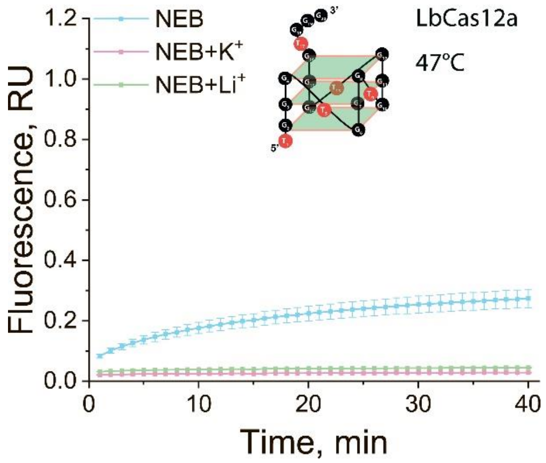
NEB
NEB+K$^+$
NEB+Li$^+$
LbCas12a
47°C
Fluorescence, RU
Time, min

A

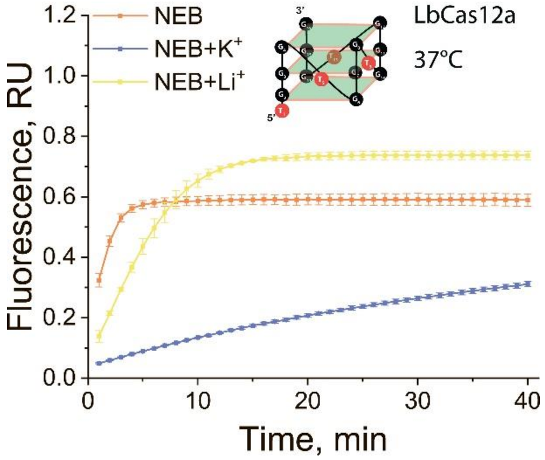
NEB
NEB+K$^+$
NEB+Li$^+$
LbCas12a
37°C
Fluorescence, RU
Time, min

B

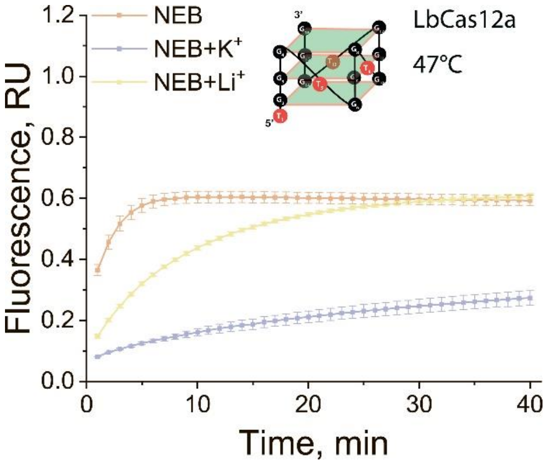
NEB
NEB+K$^+$
NEB+Li$^+$
LbCas12a
47°C
Fluorescence, RU
Time, min

C

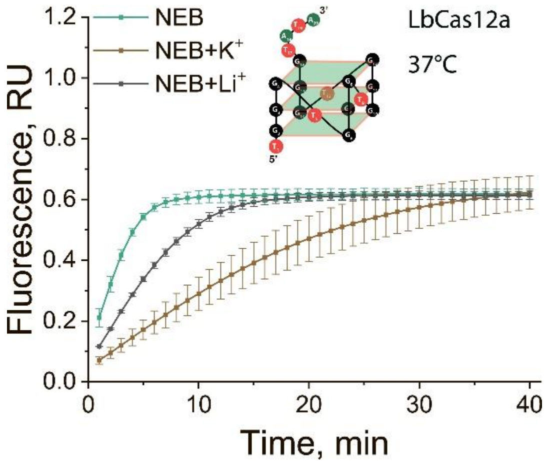
NEB
NEB+K$^+$
NEB+Li$^+$
LbCas12a
37°C
Fluorescence, RU
Time, min

D

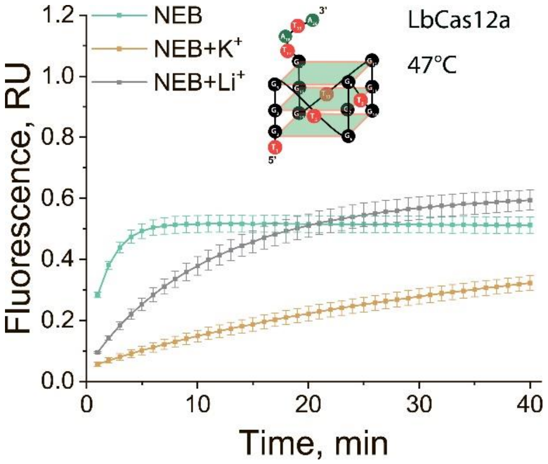
NEB
NEB+K$^+$
NEB+Li$^+$
LbCas12a
47°C
Fluorescence, RU
Time, min

E

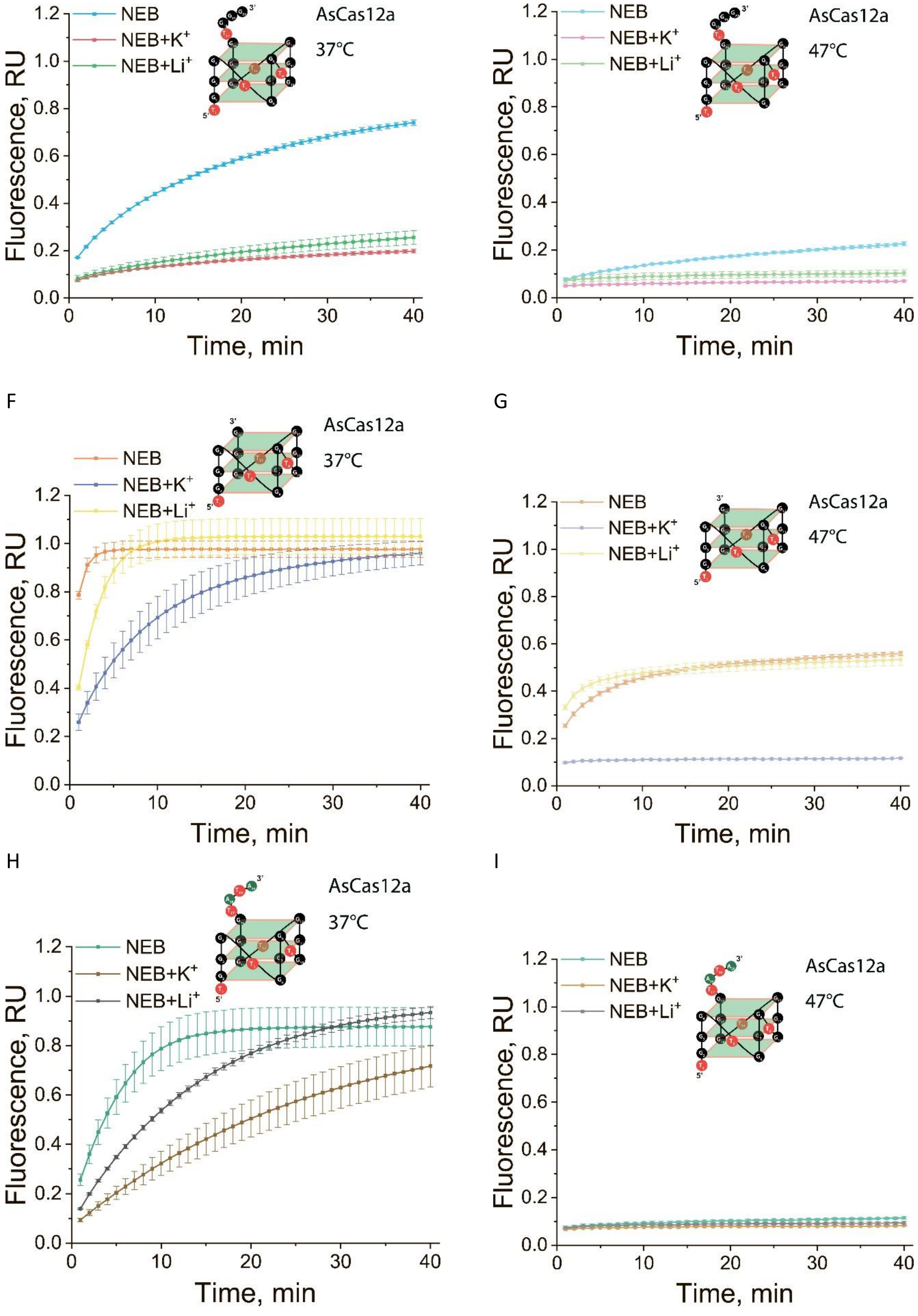


**Figure S11**. Fluorescence curves in NEB and with 50 mM KCl or 50 mM LiCl. CRISPR/LbCas12a: $(TGGG)_5$ at 47 °C (A), $(TGGG)_4$ at 37 °C (B) and 47 °C (C), $(TGGG)_4$-TATA at 37 °C (D) and 47 °C (E). CRISPR/AsCas12a: $(TGGG)_5$ at 37 °C (F) and 47 °C (G), $(TGGG)_4$ at 37 °C (H) and 47 °C (I), $(TGGG)_4$-TATA at 37 °C (J) and 47 °C (K).

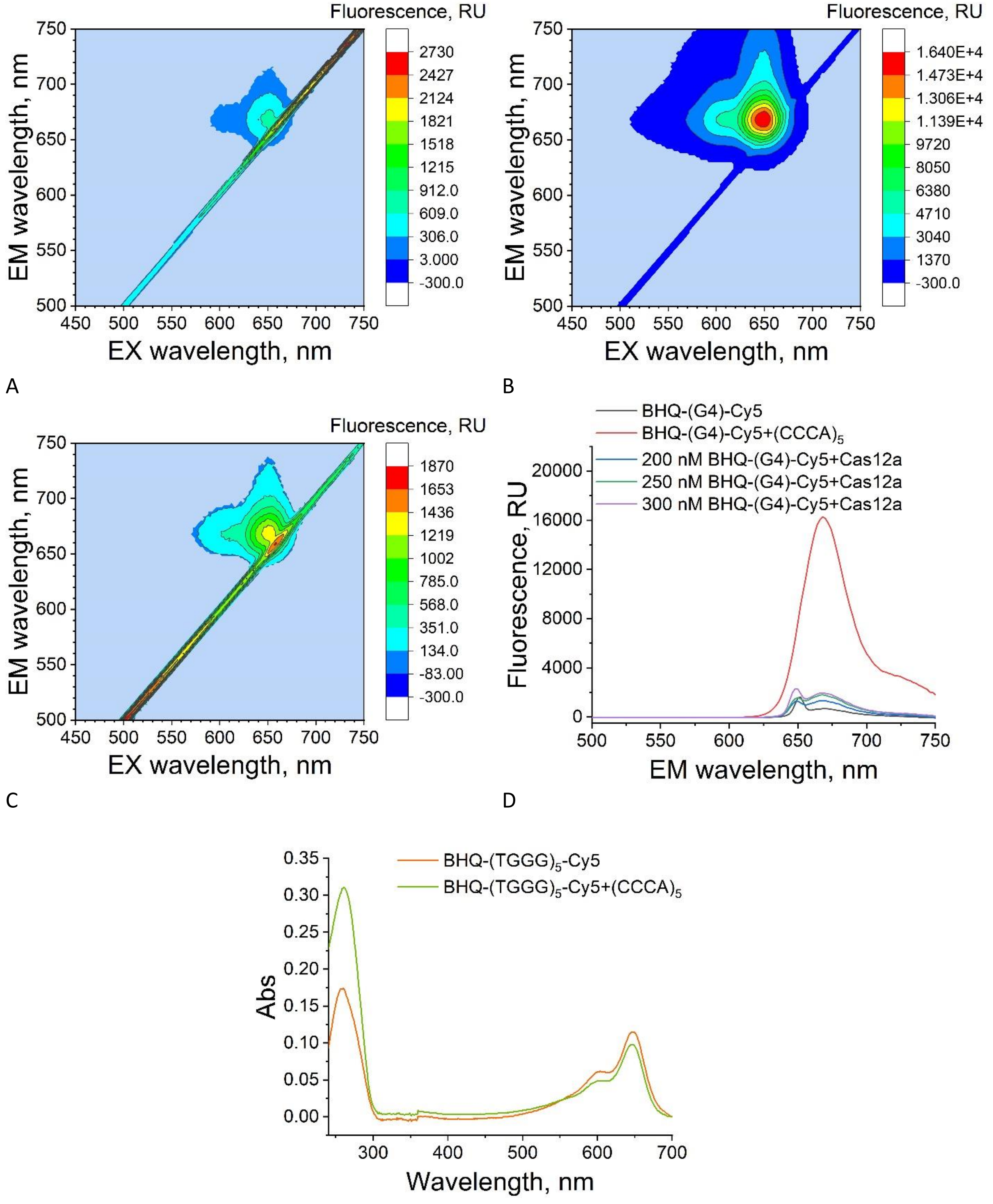


**Figure S12**. 3D fluorescence spectra (emission 500–750 nm, excitation 450–750 nm) for BHQ-(TGGG)$_5$-Cy5 (A) and the duplex BHQ-(TGGG)$_5$-Cy5 + (CCCA)$_5$ (1:2) (B), and for BHQ-(TGGG)$_5$-Cy5 after the CRISPR/Cas12a reaction (C); 2D spectra (emission 500–750 nm, excitation 648 nm) for BHQ-(TGGG)$_5$-Cy5, the +(CCCA)$_5$ duplex (1:2) and BHQ-(TGGG)$_5$-Cy5 (200/250/300 nM) after CRISPR/Cas12a (D); 2D absorbance spectra (240–700 nm) for BHQ-(TGGG)$_5$-Cy5 and the +(CCCA)$_5$ duplex (1:2) (E).

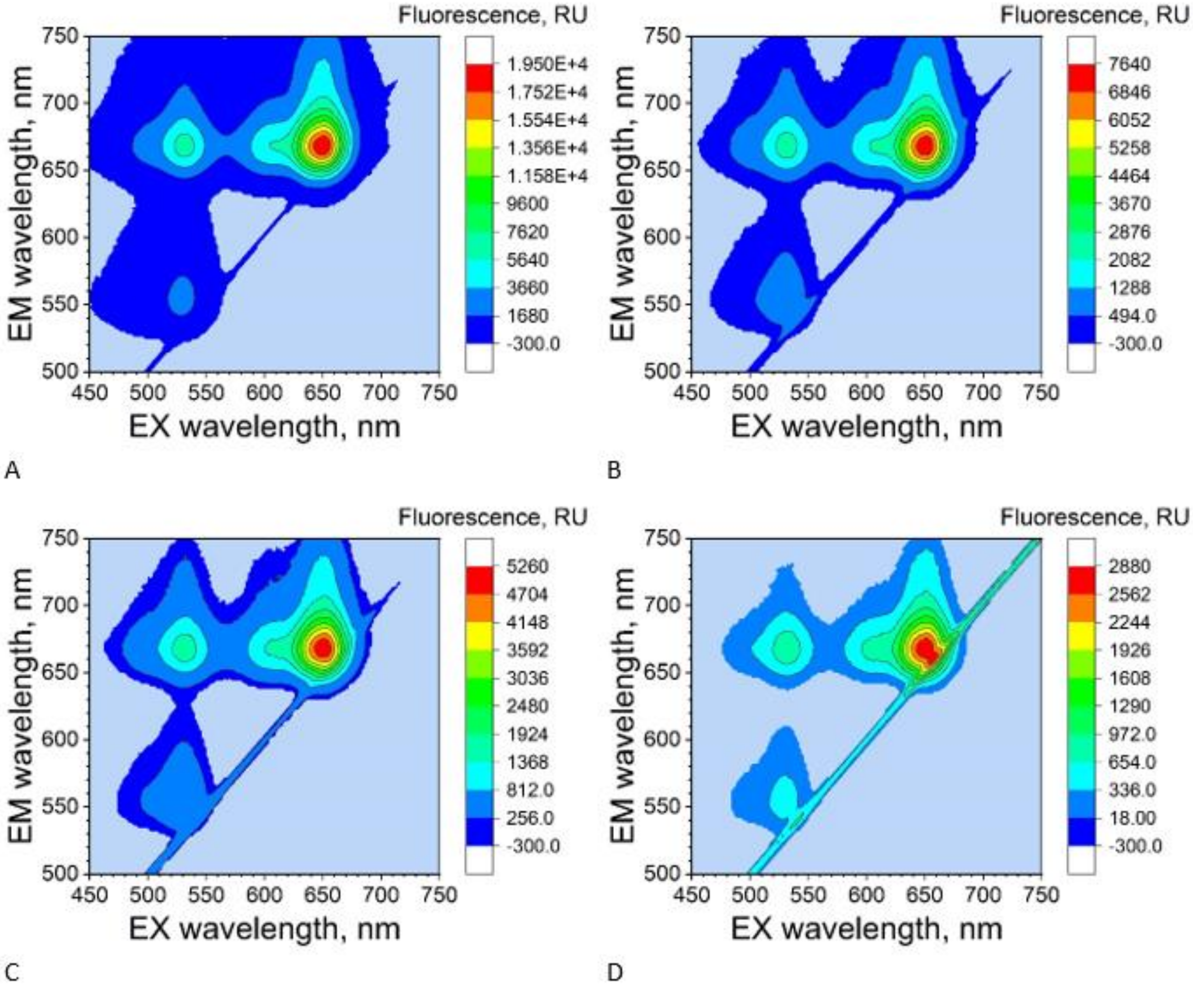


**Figure S13**. 3D fluorescence spectra (emission 500–750 nm, excitation 450–750 nm) for JOE-(TGGG)$_5$-Cy5: undiluted (A), diluted 2-fold (B), 3-fold (C) and 5-fold (D).

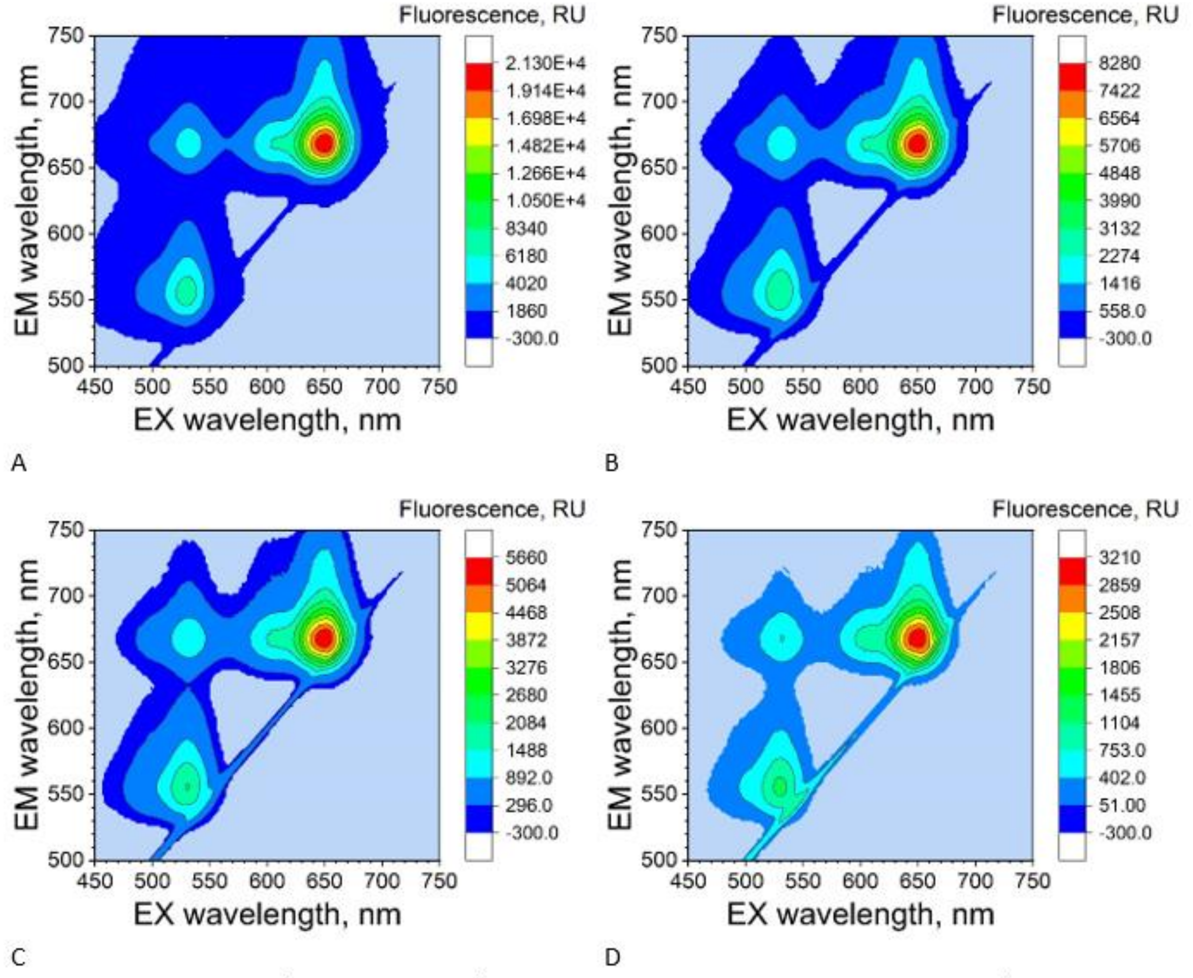


**Figure S14**. 3D fluorescence spectra (emission 500–750 nm, excitation 450–750 nm) for the duplex JOE-$(TGGG)_5$-Cy5 + $(CCCA)_5$ (1:0.5): undiluted (A), diluted 2-fold (B), 3-fold (C) and 5-fold (D).

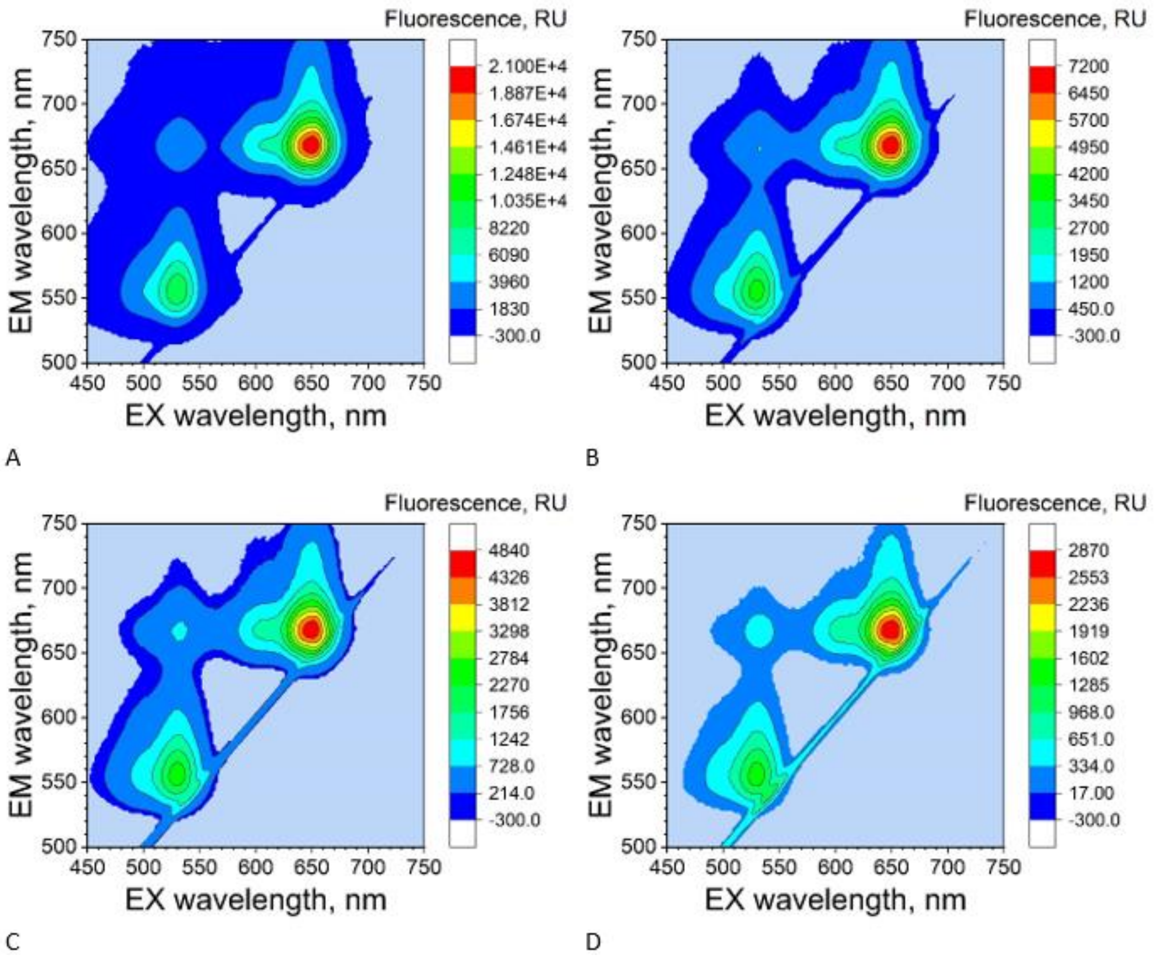


**Figure S15**. 3D fluorescence spectra (emission 500–750 nm, excitation 450–750 nm) for the duplex JOE-(TGGG)$_5$-Cy5 + (CCCA)$_5$ (1:0.75): undiluted (A), diluted 2-fold (B), 3-fold (C) and 5-fold (D).

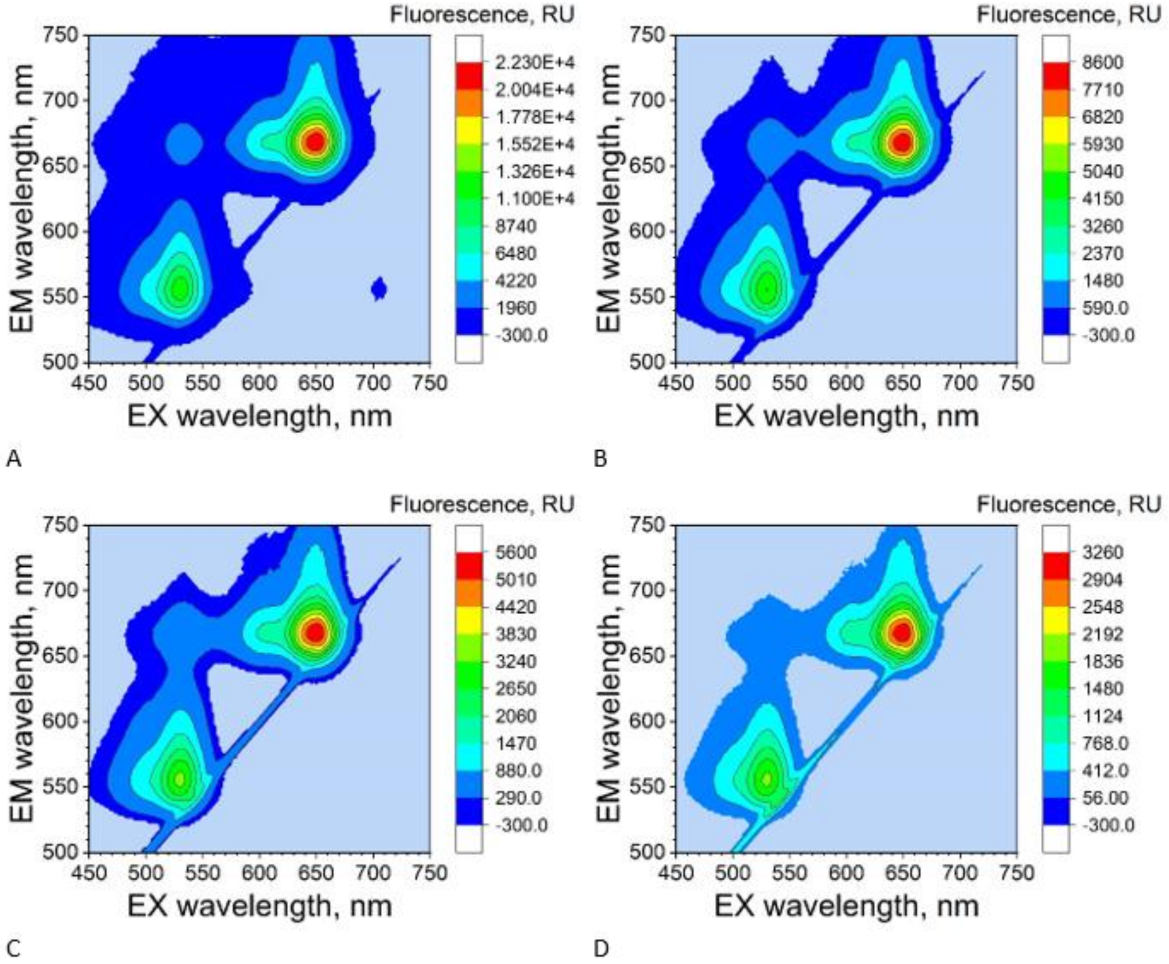


**Figure S16**. 3D fluorescence spectra (emission 500–750 nm, excitation 450–750 nm) for the duplex JOE-(TGGG)$_5$-Cy5 + (CCCA)$_5$ (1:1): undiluted (A), diluted 2-fold (B), 3-fold (C) and 5-fold (D).

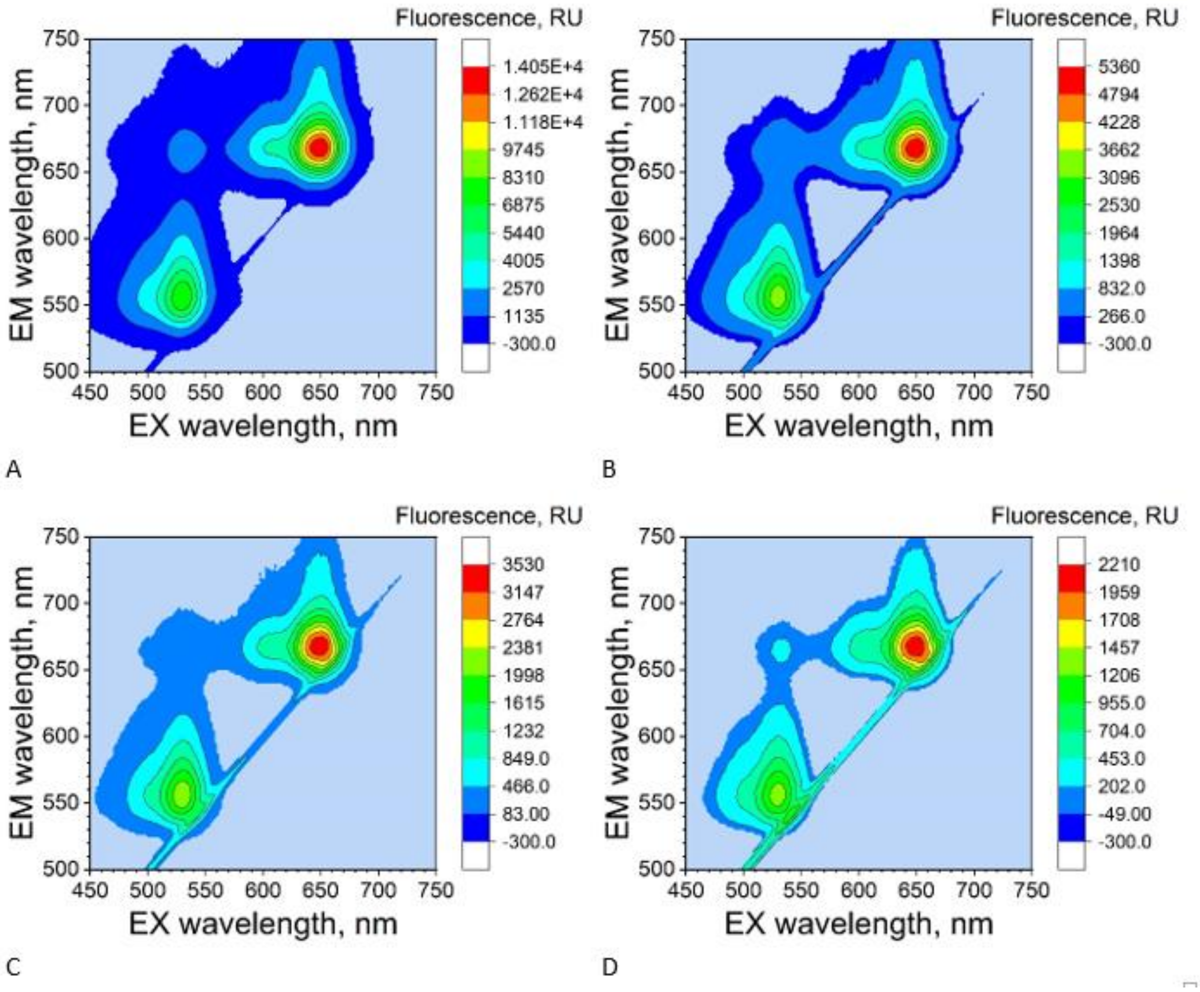


**Figure S17**. 3D fluorescence spectra (emission 500–750 nm, excitation 450–750 nm) for the duplex JOE-(TGGG)$_5$-Cy5 + (CCCA)$_5$ (1:2): undiluted (A), diluted 2-fold (B), 3-fold (C) and 5-fold (D).

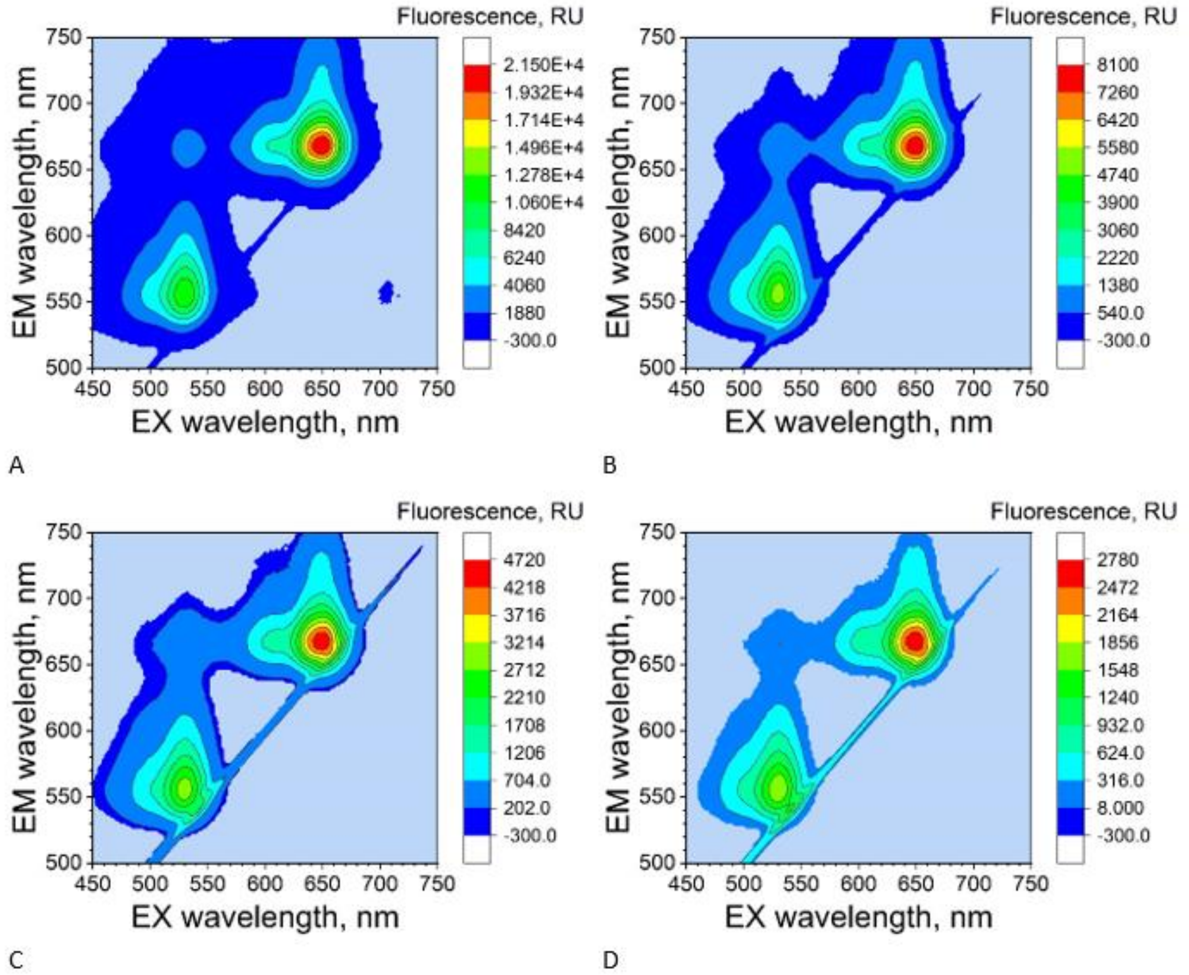


**Figure S18**. 3D fluorescence spectra (emission 500–750 nm, excitation 450–750 nm) for the duplex JOE-(TGGG)$_5$-Cy5 + (CCCA)$_5$ (1:3): undiluted (A), diluted 2-fold (B), 3-fold (C) and 5-fold (D).

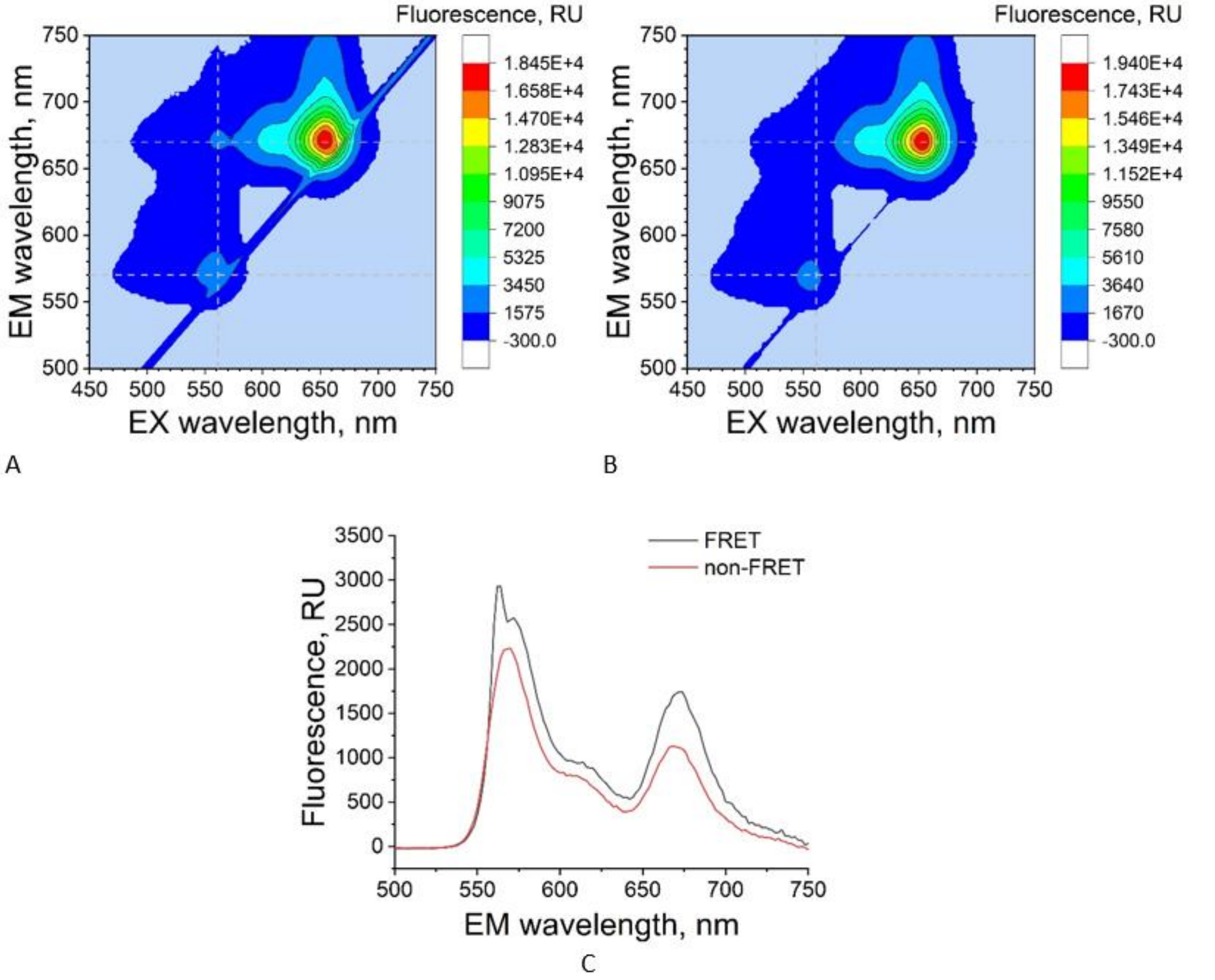


**Figure S19**. 3D fluorescence spectra (emission 500–750 nm, excitation 450–750 nm) for AF555-$(TGGG)_5$-AF647 (A) and the mixture AF555-$(TGGG)_5$-AF647 + $(CCCA)_5$ (1:2) (B); 2D spectra (emission 500–750 nm, excitation 560 nm) for AF555-$(TGGG)_5$-AF647 (FRET) and the +$(CCCA)_5$ mixture (non-FRET) (C).

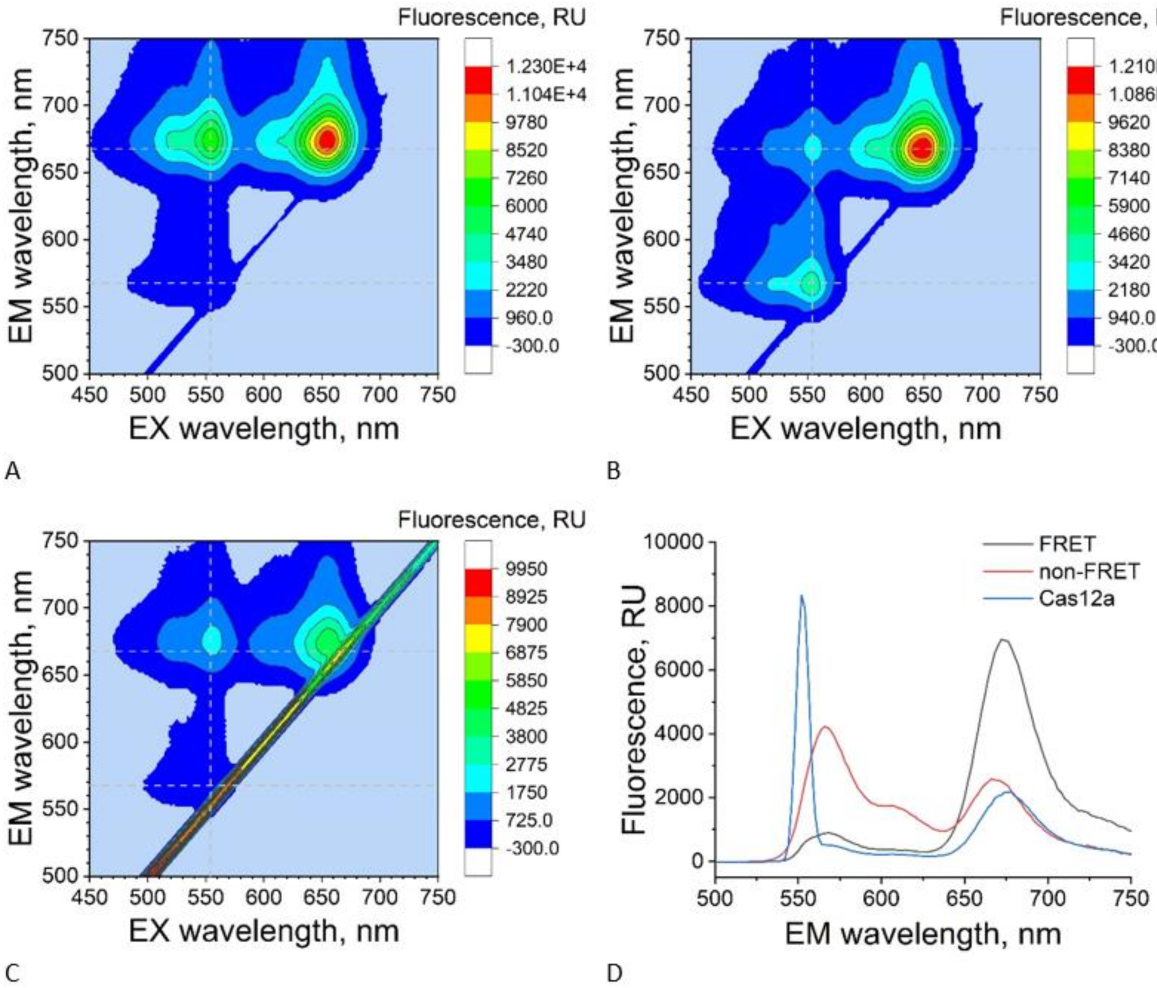


**Figure S20**. 3D fluorescence spectra (emission 500–750 nm, excitation 450–750 nm) for Cy3-$(TGGG)_5$-Cy5 (A), the mixture Cy3-$(TGGG)_5$-Cy5 + $(CCCA)_5$ (1:2) (B) and Cy3-$(TGGG)_5$-Cy5 after the CRISPR/AsCas12a reaction (C); 2D spectra (emission 500–750 nm, excitation 560 nm) for Cy3-$(TGGG)_5$-Cy5 (FRET), the +$(CCCA)_5$ mixture (non-FRET) and after CRISPR/AsCas12a (Cas12a) (D).

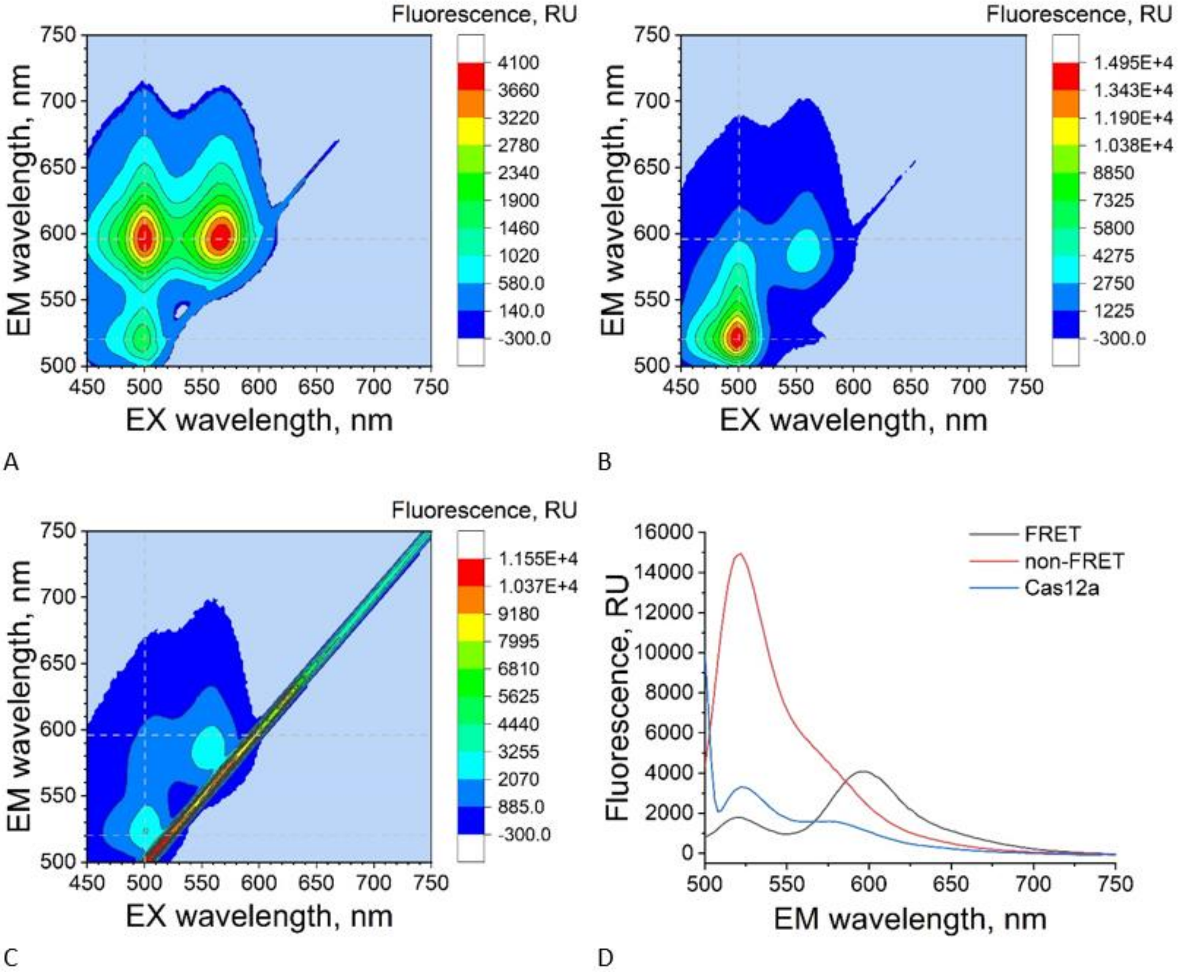


**Figure S21**. 3D fluorescence spectra (emission 500–750 nm, excitation 450–750 nm) for FAM-(TGGG)$_5$-TAMRA (A), the mixture FAM-(TGGG)$_5$-TAMRA + (CCCA)$_5$ (1:2) (B) and FAM-(TGGG)$_5$-TAMRA after the CRISPR/AsCas12a reaction (C); 2D spectra (emission 500–750 nm, excitation 560 nm) for FAM-(TGGG)$_5$-TAMRA (FRET), the +(CCCA)$_5$ mixture (non-FRET) and after CRISPR/AsCas12a (Cas12a) (D).

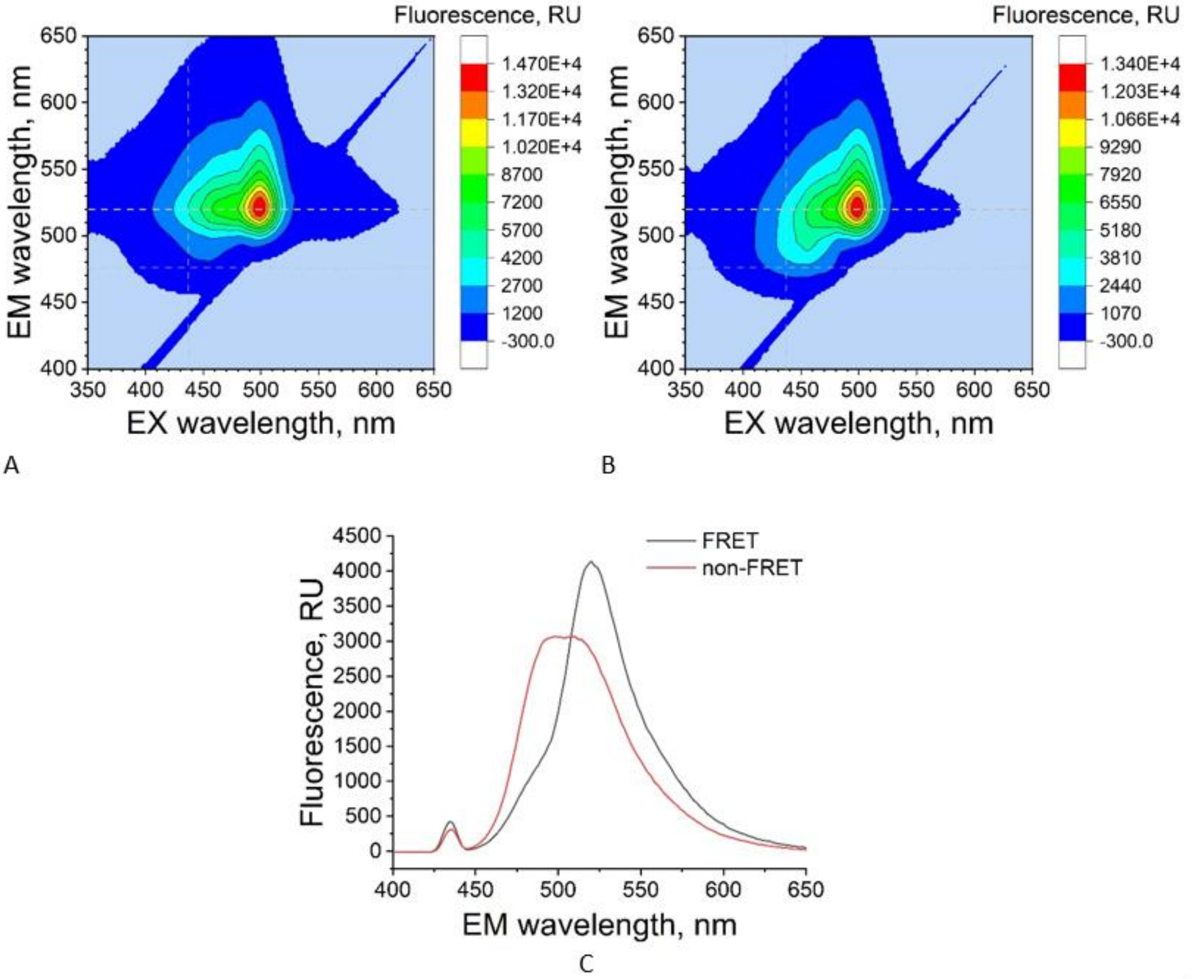


**Figure S22**. 3D fluorescence spectra (emission 500–750 nm, excitation 450–750 nm) for Coumarin343-$(TGGG)_5$-FAM (A) and the mixture Coumarin343-$(TGGG)_5$-FAM + $(CCCA)_5$ (1:2) (B); 2D spectra (emission 500–750 nm, excitation 560 nm) for Coumarin343-$(TGGG)_5$-FAM (FRET) and the +$(CCCA)_5$ mixture (non-FRET) (C).

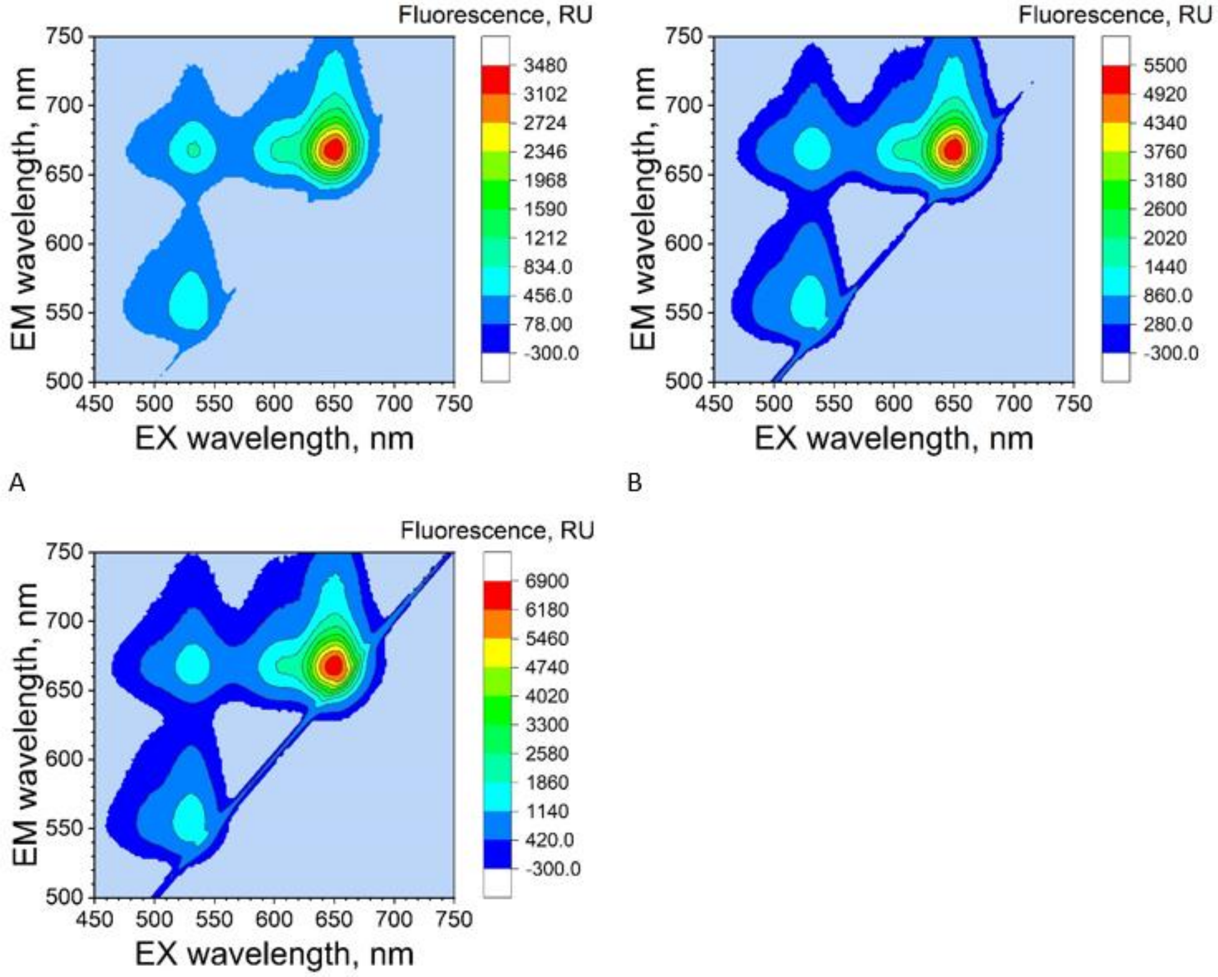


**Figure S23**. 3D fluorescence spectra (emission 500–750 nm, excitation 450–750 nm) for JOE-(TGGG)$_5$-Cy5 with gRNA-20 (1:1) at component concentrations of 200 nM (A), 250 nM (B) and 400 nM (C).

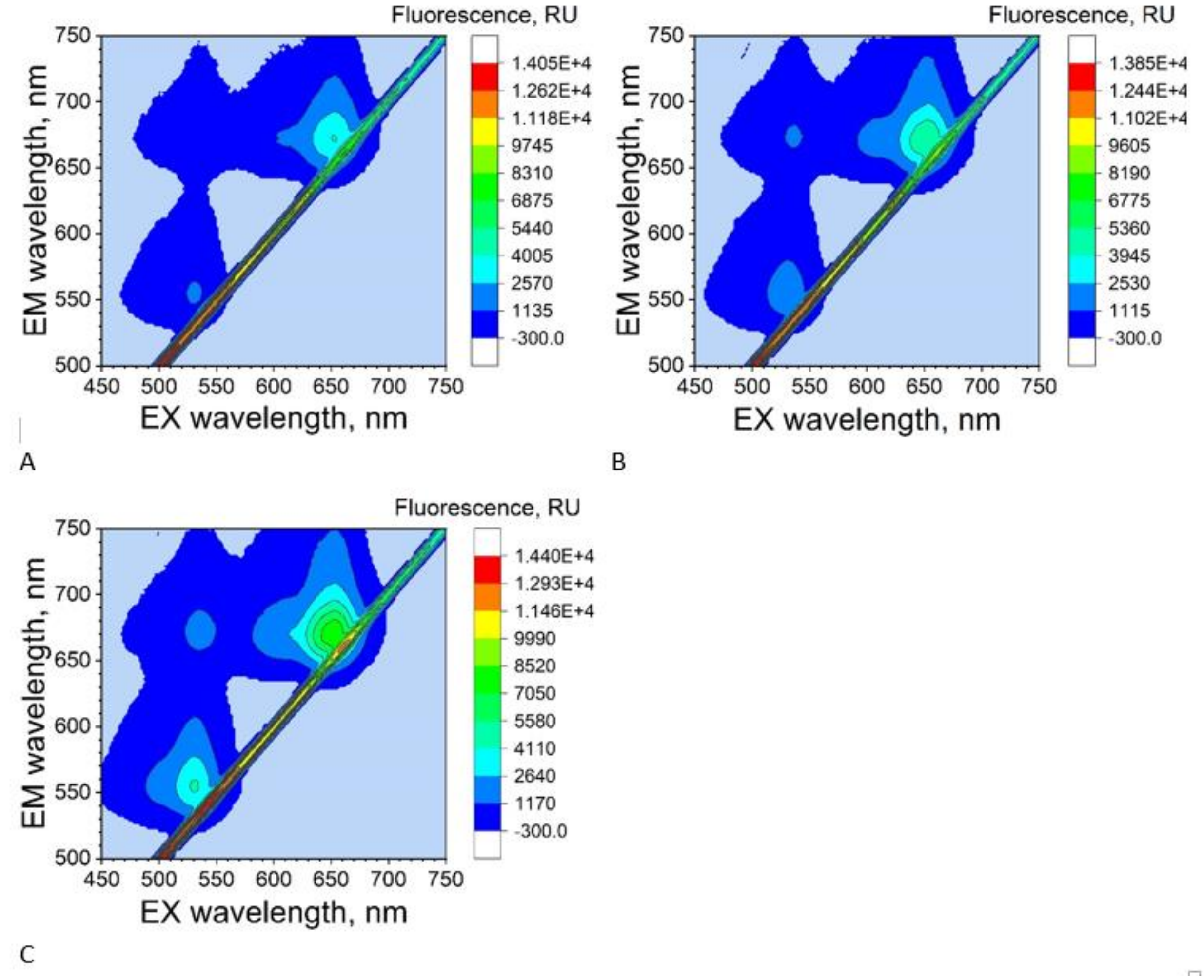


**Figure S24**. 3D fluorescence spectra (emission 500–750 nm, excitation 450–750 nm) for JOE-$(TGGG)_5$-Cy5 after the CRISPR/AsCas12a reaction with gRNA-20 (1:1) at component concentrations of 200 nM (A), 250 nM (B) and 400 nM (C).

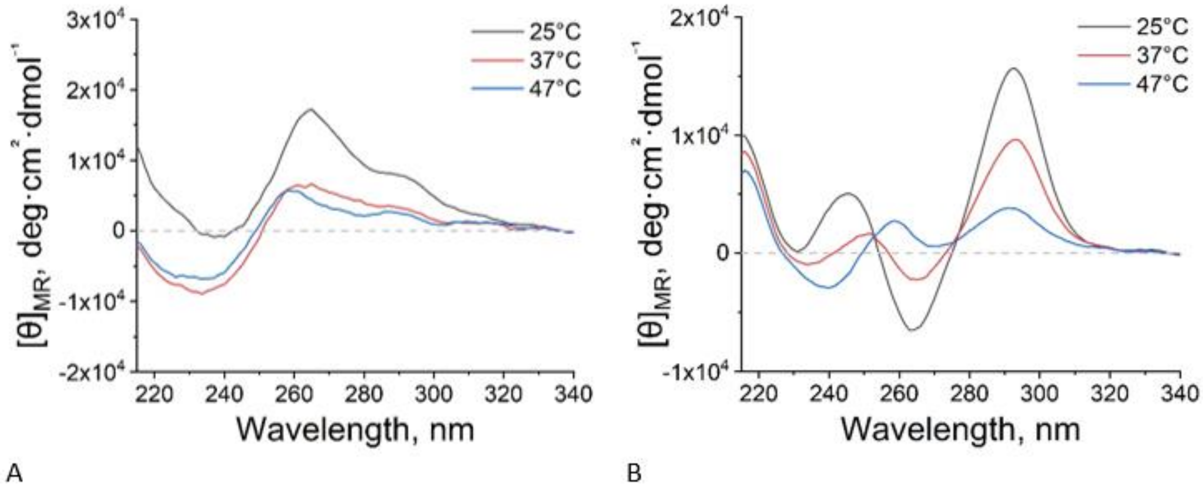


**Figure S25**. CD spectra for the antiparallel quadruplexes RE21 (A) and aptOTA (B) at 25, 37 and 47 °C in NEB buffer.

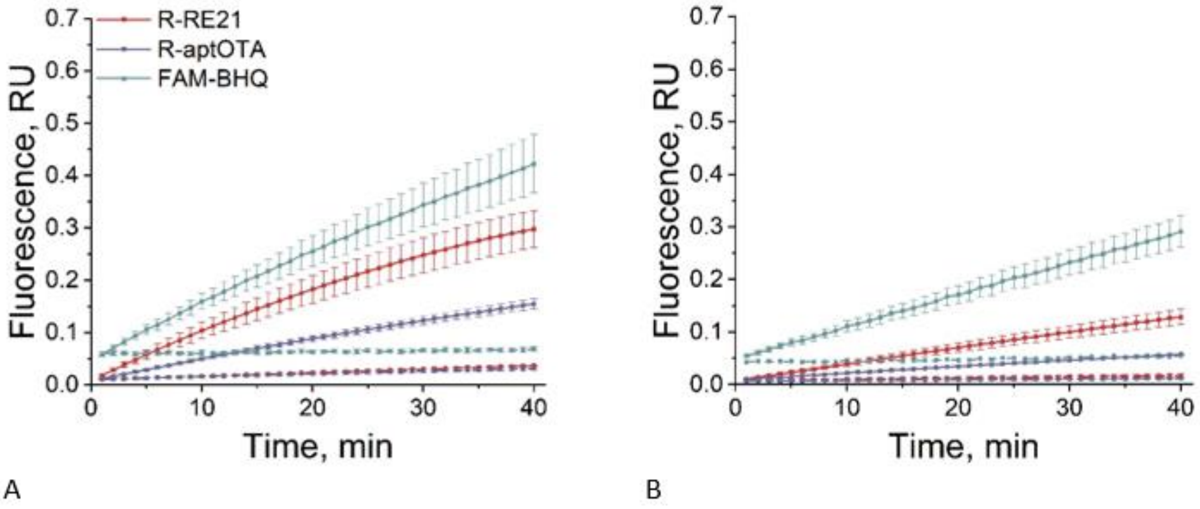


**Figure S26**. Fluorescence curves showing the role of the reporters R-RE21 and R-aptOTA as *trans*-targets in CRISPR/AsCas12a at 37 °C, compared with the standard reporter R-FAM/BHQ1, in NEB (A) and with 50 mM KCl (B); dashed lines, no-target controls (water).

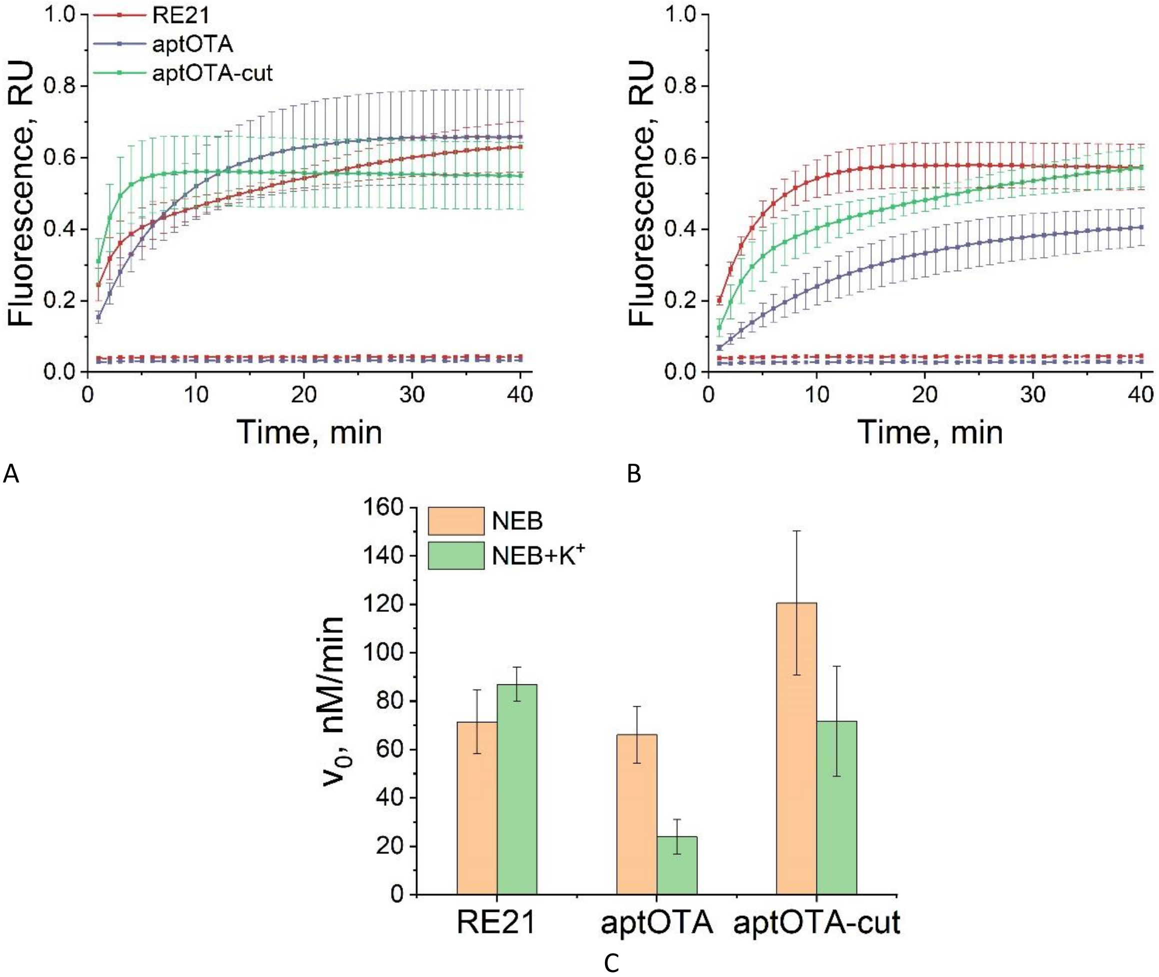


**Figure S27**. Fluorescence curves showing the role of the antiparallel G-quadruplexes RE21, aptOTA and aptOTA-cut as *cis*-targets in CRISPR/LbCas12a at 47 °C with $R(T)_{15}$ read-out, in NEB (A) and with 50 mM KCl (B); dashed lines, no-target controls (water). Comparison of initial velocities $V_0$ for RE21, aptOTA and aptOTA-cut in CRISPR/LbCas12a in NEB and with 50 mM KCl at 47 °C (C).
